# Curvature-dependent morphological reorganization of the endoplasmic reticulum determines the mode of epithelial migration

**DOI:** 10.1101/2025.04.22.650137

**Authors:** Simran Rawal, Pradeep Keshavanarayana, Diya Manoj, Purnati Khuntia, Sanak Banerjee, Basil Thurakkal, Rituraj Marwaha, Fabian Spill, Tamal Das

## Abstract

From single-cell extrusion to centimeter-sized wounds, epithelial gaps of various sizes and geometries appear in all organisms. For gap closure, epithelial cells invoke two orthogonal modes: lamellipodial crawling at the convex edge and purse-string-like movements at the concave edge. The mechanisms underlying these differential responses to geometric cues remain elusive. Here we perform an intracellular cartography to reveal that in both micropatterned and naturally arising gaps, the endoplasmic reticulum (ER) undergoes edge curvature-dependent morphological reorganizations with convex and concave edges promoting ER tubules and sheets, respectively. This reorganization depends on cytoskeleton-generated protrusive and contractile forces. Additionally, theoretical modeling predicts that the curvature-specific ER morphology leads to a lower strain energy state. ER tubules at the convex edge favor perpendicularly oriented focal adhesions, supporting lamellipodial crawling while ER sheets at the concave edge favor parallelly oriented focal adhesions, supporting purse-string-like movements. Altogether, ER emerges as a central player in cellular mechanotransduction, which orchestrates two orthogonal modes of cell migration by integrating signals from cytoskeletal networks.

---

The closure of epithelial gaps is crucial in both physiological and pathological processes. Disruptions in the epithelial barrier occur continuously across various scales and geometries, ranging from single-cell extrusions to centimeter-sized wounds, throughout an organism’s lifespan^1-6^. The epithelial gap closure happens by a collective movement of the surrounding cells^7^, which involves two orthogonal migration modes: lamellipodia-mediated cell crawling and actomyosin contractility-driven purse-string-like movements^8-12^. These modes involve different actin and focal adhesion dynamics and assembly: branched actin polymerization and perpendicularly oriented focal adhesions in lamellipodia-mediated cell crawling versus actin bundle formation and parallelly oriented focal adhesions in purse-string-like movements. Importantly, their relative contribution to epithelial gap closure depends on various biochemical and biophysical aspects of the tissue, including the curvature of the gap edge^6^. Relevantly, while previous studies have identified several molecules that sense or produce nanoscale membrane curvatures^13, 14^, how cells coordinate a response toward a curvature of cellular and multicellular length-scale remains largely elusive. Nevertheless, it is known that the epithelial cells prefer to undergo lamellipodia-mediated cell crawling at the positively-curved convex edge and purse-string closure at the negatively-curved concave edge^15^. Concomitantly, the speed of the migrating cells depends upon the gap geometry. Also, a previous study has shown how large-scale curvature dependence is regulated by directional actin flow, where the actin shows anterograde flow at the concave edge as opposed to retrograde flow during lamellipodia formation at the convex edge^16^. However, it remains unknown how other intracellular structures organize in response to large-scale curvature, how such organizations are regulated, and whether they influence the mode of cell migration.

Here, we used both micropatterned wounds of defined geometry in cultured epithelial monolayer and natural wounds with spontaneously arising convex and concave curved regions in mouse embryonic epidermis and performed the cartography of intracellular structures. We imaged the structure and organization of the cytoskeletal elements and membrane-bound cell organelles and discovered that the endoplasmic reticulum (ER) shows a drastic morphological rearrangement in response to curvature. ER is a continuous and highly dynamic organelle that spans the whole cell as a single entity. It comprises rounded tubular networks and dense flat sheets, which undergo constant rearrangements between themselves^17-23^. This highly dynamic nature of the ER reflects its adaptability and responsiveness to intracellular and extracellular changes^24-26^. However, the relationship between the dynamic ER morphologies and physical or geometric cues remains largely unknown. Here we report that the endoplasmic reticulum alters its morphology and dynamics in response to edge curvature and plays a critical role in determining the modes of epithelial migration at different edge curvatures.

## RESULTS

### Edge curvature-dependent ER morphology in micropatterned gaps

First, to reveal the cell biological mechanism underlying the differential response of epithelial cells to edge curvature, we used microfabrication^8,15,16^, which provided a controlled and reproducible means to generate gaps of a definite shape, size, and edge curvature. We fabricated polydimethylsiloxane (PDMS) stencils of various shapes consisting of convex, concave, and flat edges^15^. To study the collective migration of the epithelial cells at different edge curvatures, we seeded Madine-Darby canine kidney (MDCK) epithelial cells around these PDMS stencils (Fig. 1a) for 15-18 hours and allowed them to form a confluent monolayer. Subsequently, we lifted off the stencils triggering the cells to start migrating into the gaps. Considering that the lamellipodial protrusions or actin bundles emerge within 15-30 minutes^15^, we fixed the cells 30 minutes after removal of the stencil and stained them with fluorescent dye-labeled phalloidin to visualize the organization of the actin cytoskeleton. As expected^15^, we observed that the fraction of cells forming lamellipodia at the positively-curved convex edge was significantly higher than that at the negatively-curved concave edge (Fig. 1b, upper panel, Supplementary Fig. 1a). We next studied the arrangement of the other major cytoskeletal element, the microtubules. Like actin, microtubules also appeared to have differential arrangements at convex and concave edges (Fig 1b, middle panel). Microtubule filaments appeared more discrete and aligned perpendicular to the edge at the convex edge while they formed thick bundles aligned parallel to the edge at the concave edge (Fig. 1b, middle panel). Quantification of the microtubule directionality further affirmed this conclusion (Supplementary Figs. 1b-c).

**Figure 1.**
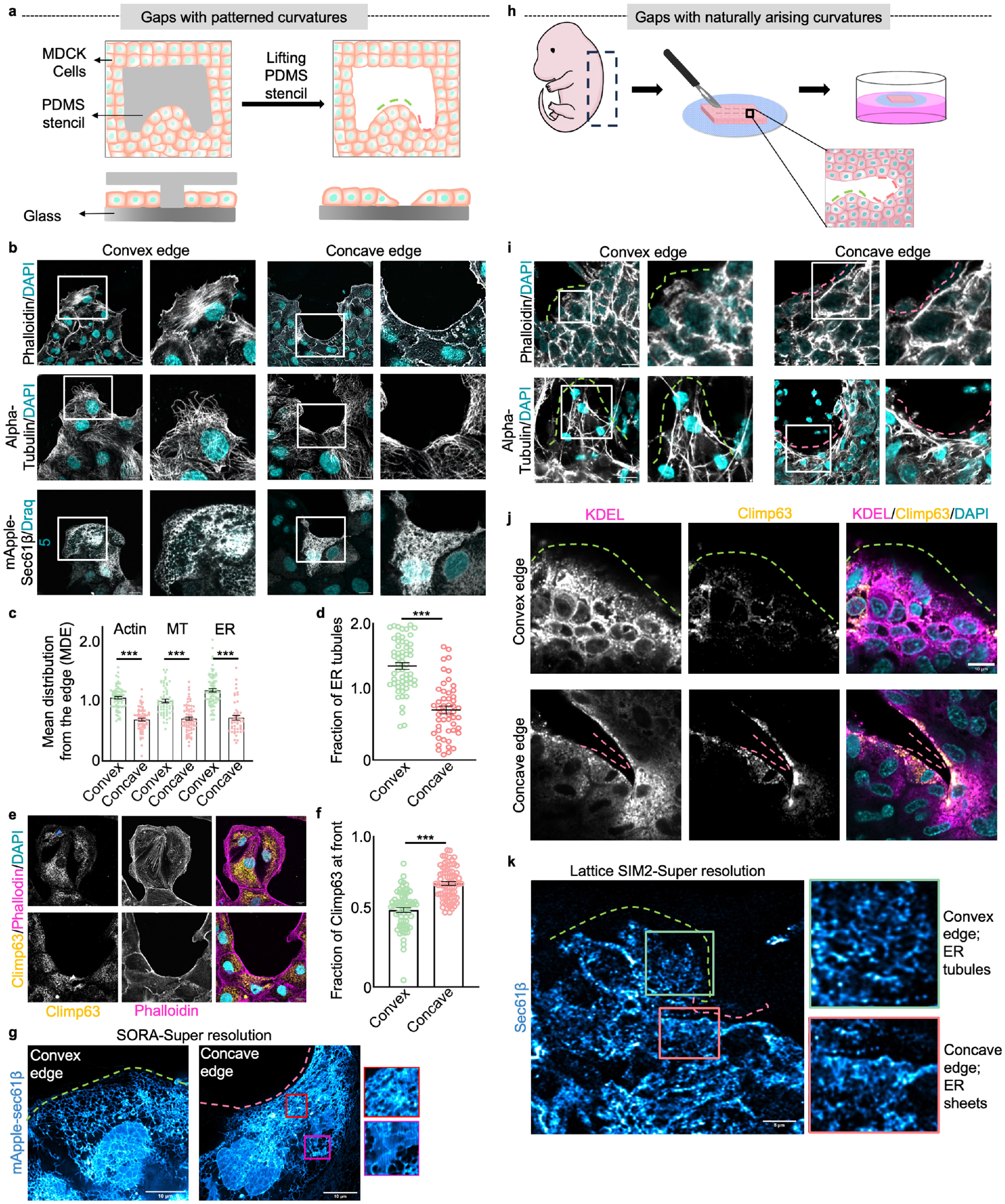
Edge curvature-dependent ER morphologies. **a**, Schematic representing the experimental setup for creating gaps of definite geometries. Cells surround the PDMS stencils placed on glass, after removal of which cells migrate into the voids. **b**, Representative images of actin, microtubule, and ER at convex (left) and concave (right) edges. MDCK cells stained with top panel-phalloidin (grey, Actin marker) and DAPI (cyan) ; middle panel-anti-α-tubulin (grey, microtubule marker) and DAPI (cyan); bottom panel - MDCK cells expressing mApple-Sec61β (grey, ER marker) and labeled with DRAQ5 (cyan). Edge of the cells near the curvature enlarged on the right of individual images, scale bar - 10μm **c**, MDE quantifications for ER, actin, and microtubules at convex (green) and concave edges (pink). n= 74,62,62,69,83,49 from left to right **d**, quantification for fraction of ER tubules present at the front of the cells at the convex (green) and concave (pink) edge n=58,50 from left to right **e**, Representative images of cells stained with anti-Climp63 (yellow, ER sheet marker), phalloidin (pink) and DAPI (cyan) at convex (upper panel) and concave edge (lower panel), blue arrowhead shows low Climp63 at the edge of cell, white arrowhead shows Climp63 enrichment at the edge, scale bar: 10μm. **f**, quantification of Climp63 intensity at front of the cells at convex (blue) and concave (pink) edge n=68, 71 from left to right **g**, Representative super-resolution images of MDCK cells expressing mApple-Sec61β at convex (left) and concave (right) edge, insets of the concave edge show, fenestered sheets (red inset) and flats sheets (magenta inset). **h**, Schematic representing *ex vivo* wound healing experiment **i**, Representative images of actin and microtubule at convex (left) and concave (right) edge in mouse embryonic skin wounds. E14.5 wounded skin stained for the upper panel-phalloidin (grey) and DAPI (cyan); lower panel-anti-α-tubulin (grey) and DAPI (cyan). Edge of the cells near the curvature enlarged on the right of individual images, scale bar: 10μm. **j**, Representative images of wounded embryonic skin stained with anti-KDEL (magenta, ER marker), anti-Climp63 (yellow) and DAPI (cyan) at convex (upper panel) and concave (lower panel) edges, Edge of the tissue marked with dashed lines, scale bar: 10μm. **k**, Super-resolution images of wounded embryonic skin stained for anti-Sec61β antibody, convex (green inset) and concave (pink inset) edges zoomed and enlarged at the right, scale bar 5μm. Data are mean ± s.e.m., Anova test **(c)**, Two-tailed t-test **(d**,**f)** Pink dashed lines represent a concave edge, and green dashed lines represent a convex edge.

Next, to investigate whether other intracellular entities also respond to this large-scale geometrical cue, we performed a cartographic study of intracellular organization at differently curved edges, focusing on cellular organelles. We used anti-GRASP65 antibody, anti-LAMP1 antibody, and mitotracker green to visualize the Golgi apparatus, lysosomes, and mitochondria, respectively. In addition, we transfected MDCK cells with mApple-Sec61β to observe the ER structure. Golgi showed a ribbon-like structure at both curvatures, but it was mostly polarized in the front of the nucleus at the convex edge and next to the nucleus along its horizontal axis at the concave edge (Supplementary Fig. 1d). Lysosomes appeared closely localized near the actin bundle at the concave edge, but more dispersed in the lamellipodia at the convex edge (Supplementary Fig. 1d). Mitochondria had filamentous morphology spread throughout the cell and showed no apparent difference in structure or organization depending on the edge curvature (Supplementary Fig. 1d). Interestingly, ER morphology appeared significantly different within the cells located at convex and concave edges. We observed the formation of a reticular network of ER tubules at the convex edge and a dense sheet-like lamellar morphology at the concave edge (Fig. 1b, lower panel). Beyond qualitative visualization, we also quantitatively characterized the distribution of cytoskeleton and organelles at different edge edges by using a statistical approach^27^, which is based on probability density estimation. In this respect, we calculated the mean distribution from the edge (MDE), which quantifies the distance between the edge of the cell and the center of mass of the intracellular entity (Supplementary Fig. 1e). A lower value of the MDE suggests that the organelle or the cytoskeleton is polarized and densely accumulated near the migrating edge. We observed that the MDE for actin, microtubules, ER, Golgi apparatus, and lysosomes at the concave edge is significantly lower than the respective MDE at the convex edge suggesting that they are differentially polarized in response to the geometrical cue (Fig. 1c, Supplementary Fig. 1f). MDEs of mitochondria and nucleus did not show any significant difference at convex and concave edge (Supplementary Fig. 1f). Taken together, of all organelles, ER showed differences in not just the distribution but also in its morphology and structure at different edge curvatures.

We quantified the structural changes in ER by calculating the fraction of ER tubules at the front of the cell by segmenting the ER into tubules and sheets using trainable Weka segmentation^28^. This analysis revealed a high ER tubule fraction at the front of cells at the convex edge as compared to in cells at the concave edge (Fig. 1d). Next, to confirm whether the dense ER at the front of migrating cells at the concave edge are indeed ER sheets, we labeled the cells with an antibody against cytoskeleton-linking membrane protein 63 (Climp63), which is an ER sheet marker. We observed that at the convex edge, Climp63 localized mostly around the nucleus at the cell center, whereas at the concave edge, it accumulated at the cell periphery facing the edge (Figs. 1e-f). To further reveal the edge curvature-dependent ER morphology at a higher resolution, we finally performed a super-resolution imaging of cells transfected with mApple-Sec61β using super-resolution by optical realignment (SoRa) at different edge curvatures. We observed that a highly reticulated network of ER tubules was spread at the convex edge. In contrast, at the concave edge, we observed accumulation of the dense ER which consisted of flat ER sheets and fenestrated ER sheets (Fig. 1g). We also verified the generality of these observations in another epithelial cell line, namely Eph4-Ev, which originated from the mouse mammary gland (Supplementary Fig. 2).

### Edge curvature-dependent ER morphology in mouse embryonic epidermis

Next, we examined the cellular response to spontaneously arising edge curvature in an incision wound made in mouse embryonic skin explant (Fig. 1h). In this *ex vivo* model, we harvested the dorsal skin explants from E14.5 to E16.5 mouse embryos, created incisional wounds in the epidermis, and maintained them in an air-liquid interface to allow migration. Then after 30 minutes, we fixed the tissues and stained them for actin, microtubules, and ER. Phalloidin staining depicted that the epidermal cells formed lamellipodia at the convex wound edge and supracellular actin bundles at the concave wound edge (Fig. 1i, Supplementary Fig. 3a). Microtubule orientation also depended on the local wound curvatures. They showed perpendicular and parallel alignments at the convex and concave wound edges, respectively (Fig. 1i, Supplementary Fig. 3b). Finally, to study ER morphology in the mouse embryonic skin wounds, we stained the skin explants with an anti-Climp63 antibody. Very similar to the results from the micropatterned edges (Figs. 1e-f), Climp63 was specifically localized at the cell periphery at the concave wound edge. In contrast, staining these explants with antibody against the KDEL sequence, a general ER marker, revealed that ER network covered almost the entire cytoplasm (Fig. 1j). Together, these results indicated specific enrichment of ER sheets at the concave wound edge. To resolve the structural details of ER in the mouse skin tissue further, we used Lattice SIM^2^ super-resolution imaging and observed that ER indeed depicted more tubular morphology at the spontaneously generated convex wound edge and dense sheet-like structures at the concave wound edge (Fig. 1k). Taken together, these observations suggested that epithelial cells interfacing different edge curvatures have proclivity for generating different ER morphologies.

### Differential ER dynamics at convex and concave edges

To understand how distinct ER structures might be linked to ER dynamics at different edge curvatures, we performed live cell imaging using MDCK cells expressing LifeAct, which marked the actin cytoskeleton, and mApple-Sec61β, which marked the ER. As mentioned above, we seeded these cells around the micropatterned pillars, incubated for 15-18 hours, and subsequently, lifted off the stencils allowing the cells to start migrating into the gaps. Live imaging revealed two kinds of ER dynamics at the front of the migrating cell. At the convex edge, we observed a retrograde flow of dense ER structures, towards the nucleus where major ER sheets are present (Fig. 2a, Supplementary video 1). Plotting a kymograph affirmed this retrograde flow (Fig. 2b). In contrast, at the concave edge, we observed an anterograde movement of ER sheets towards the front of the migrating cells (Fig. 2c, Supplementary video 2), as also affirmed by the kymograph (Fig. 2d). A high-resolution imaging further revealed that at the convex edge, as the cells started migrating, newer ER tubules grew in the direction of migration (Fig. 2e, green arrowhead, Supplementary video 3) and also, existing tubules retracted back into the tubular ER network (Fig. 2e, red arrowhead, Supplementary video 3). In contrast, at the concave edge, the tubules grew towards the cell periphery, accumulated there, and formed sheets (Fig. 2e, Supplementary video 4). Interestingly, we also observed anterograde movements of the existing sheets towards the edge. Subsequently, we quantified the fraction of ER tubules at the cellular front at different time points. This analysis revealed that a bias existed at the beginning where ER tubule fraction was determined to be 0.64 and 0.35 at the convex and concave edges (Fig. 2f). As time progressed, the fraction of tubules at the convex edge increased further, while it decreased at the concave edge (Fig. 2f). Taken together, these results suggested that contrasting ER morphologies at the two edge curvatures are coupled with differential dynamics of ER.

**Figure 2.**
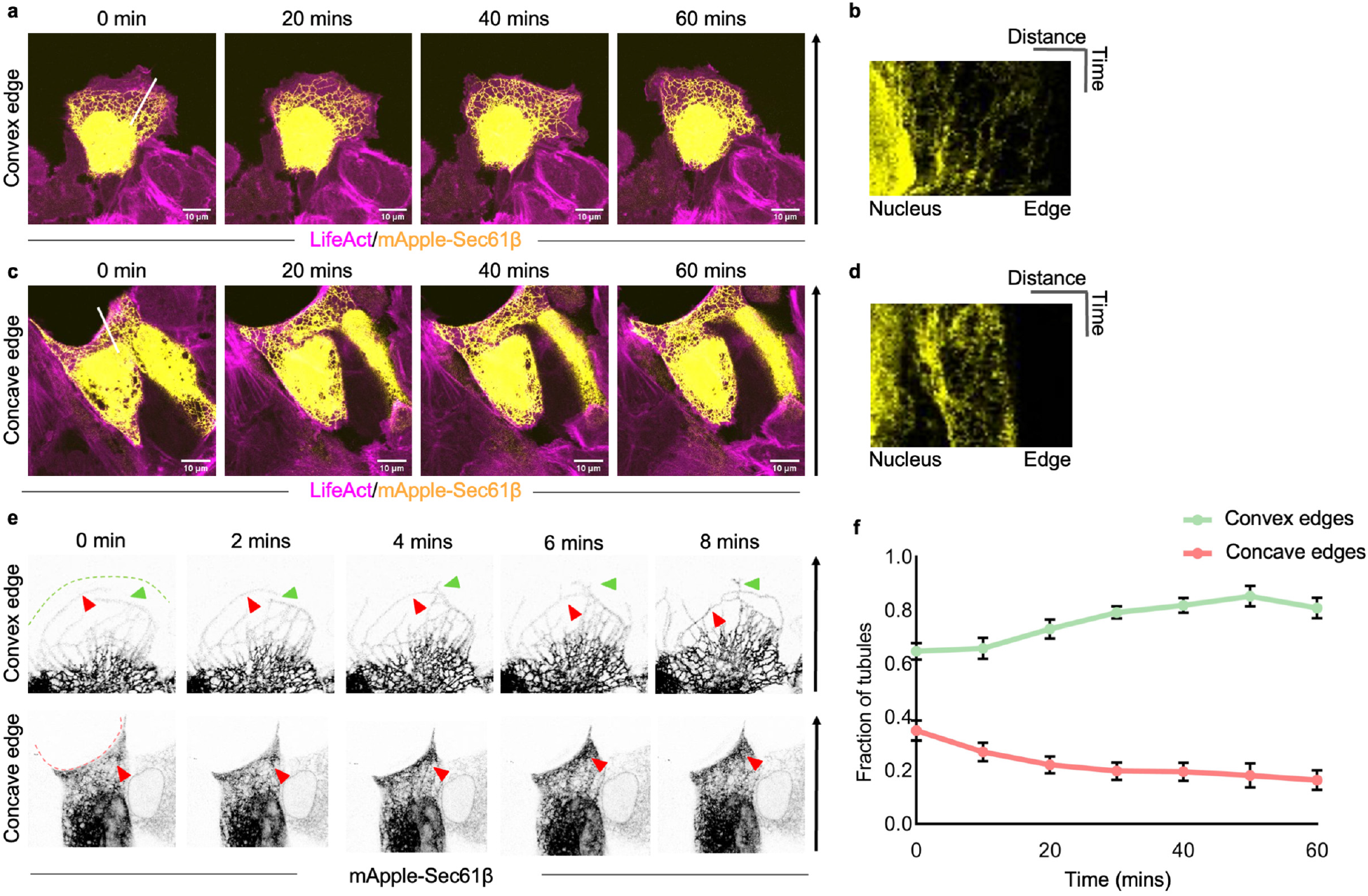
Differential ER dynamics at convex and concave edges. **a**, Time lapse images of LifeAct MDCK cells expressing mApple-Sec61β migrating at convex edge scale bar-10μm **b**, Kymograph of the line in **a** representing movement of the ER scale bar-5μm, 20mins **c**, Time lapse images of LifeAct MDCK cells expressing mApple-Sec61β migrating at concave edge scale bar-10μm **d**, Kymograph of the line in **c** representing movement of the ER scale bar 5μm, 20mins **e**, Timelapse images of front of MDCK cells expressing mApple-Sec61β at convex (upper panel) and concave (lower panel) edge. Green arrowheads show ER tubule growing in the direction of migration, red arrowhead (upper panel) shows retrograde movement of the ER tubule, red arrowhead (lower panel) show anterograde movements of tubules towards the edge **f**, quantification of ER tubule fraction in the cells over 60 mins at the two curvatures (n=10 at each curve). Black arrow represent direction of migration (**a**,**c**,**e**).

### Protrusive and contractile forces regulating ER morphology at different edge curvatures

Epithelial cells exposed to the wound respond by generating protrusive forces due to branched actin polymerization at the convex edge and contractile forces due to actomyosin activity at the concave edge^15,16,29^. We investigated whether the microtubules and ER responded to these forces by altering their orientation and morphology, respectively. We first treated the cells with an Arp2/3 inhibitor, CK666, which prevented lamellipodia formation, attenuating the protrusive force. CK666-treated cells predominantly formed actin bundles at both convex and concave edges (Supplementary Figs. 4a-b). Importantly, in these cells, at both edges, microtubules were arranged as parallel bundles (Figs. 3a-b). Simultaneously, ER showed sheet morphology at the edge of cells (Figs. 3c-d), as opposed to the control cells where bundled microtubules and ER sheets were prevalent mainly at the concave edge. We next treated the cells with Blebbistatin, inhibiting the actomyosin contractility. Blebbistatin-treated cells formed protrusive structures at both convex and concave edges (Supplementary Figs. 4a-b). In these cells, microtubules aligned perpendicularly to the edge (Figs. 3a-b), and ER showed tubular structure at both convex and concave edges (Figs. 3c-d). We, however, wanted to confirm whether CK666 or blebbistatin treatments affected ER structure globally and therefore, examined several cells in the middle of the monolayer. Overall morphology of ER was not affected, rather the effect was confined at the edge of the monolayer (Supplementary Figs. 4c-d). These results suggested microtubules and ER responded to different kinds of forces generated by the protrusive and contractile actin structures at different edge curvatures.

**Figure 3.**
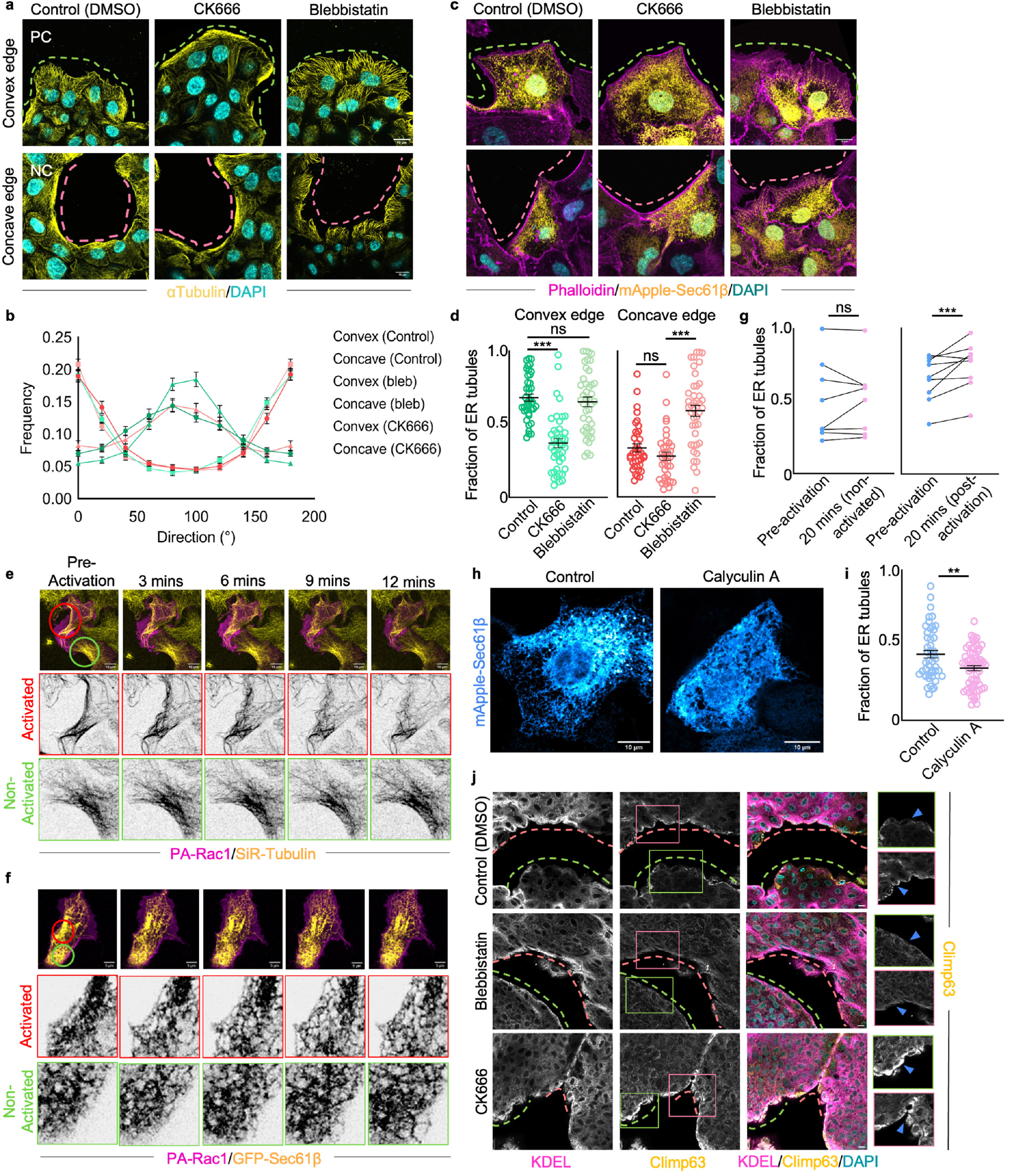
Protrusive and contractile forces regulate ER morphologies at two curvatures. **a**, Representative images of MDCK cells stained with anti α-tubulin (Yellow) and DAPI (cyan) treated with DMSO (left), CK666 (middle) and Blebbistatin (right) migrating at convex (upper panel) and concave edge (lower panel) scale bar-10um **b**, quantification for directionality of microtubules in cells treated with CK666 or Blebbistatin at different curvatures (n=75) **c**, Representative images of MDCK cells expressing Sec61β (yellow) stained with phalloidin (magenta) and DAPI (cyan) treated with DMSO (left), CK666 (middle) and Blebbistatin (right) at convex (upper panel) and concave (lower panel) edge scale bar-10μm **d**, quantifications for fraction of tubules at the front of the cells treated with DMSO, CK666, or Blebbistatin at convex (green, left) and concave (pink, right) edge n= 40, 42, 41 from left to right **e**, Time lapse imaging of cells expressing PA-Rac1 (magenta) labelled with SiR-tubulin (yellow) photo-activated with 445nm laser in region shown with the red circle, green circle represents a part of non-activated region, zoomed in regions show SiR-tubulin at activated and non-activated regions scale bar - 10μm **f**, Time lapse images of cells expressing PA-Rac1 (magenta) and GFP-Sec61β (yellow), photo-activated with 445nm laser in region marked with red circle, green circle represents non activated region, zoomed images of GFP-sec61β at activated and non-activated regions shown scale bar-5μm **g**, quantification of fraction of ER tubules in photo activated regions before and after 20mins of activation (right) and non-activated regions (left) before and after 20 mins, n=10 each **h**, representative images of MDCK cells expressing mApple-Sec61β treated with DMSO (left) and Calyculin A (right) scale bar-10μm **i**, quantification of the fraction of tubules in cells treated with DMSO or calyculinA n= 49, 66 from left to right **j**, representative images of mouse embryonic skin wounds stained with KDEL (magenta), Climp63 (yellow), and DAPI (cyan) and treated with DMSO (Top panel), Blebbistatin (middle panel) and CK666 (bottom panel), Green insets show Climp63 in cells zoomed in at convex edge, Pink inset show Climp63 in cells zoomed in at concave edge, blue arrowheads point towards the edge of the cells, scale bars, 10 μm. Data are mean ± s.e.m., Anova test **(d)**, Two tailed t-test **(i).** Pink dashed lines mark the concave edges, green dashed lines mark the convex edges **(a**,**c**,**j)**

Next, to perturb the system in a more controlled manner, we used optogenetic tools to drive lamellipodia formation and followed the morphological changes in ER and microtubules using live cell imaging. Rac1 is a small Rho GTPase that regulates actin assembly and dynamics, especially during lamellipodia formation. We used a genetically encoded photoactivatable Rac1 (PA-Rac1) which induces the formation of lamellipodial protrusions on activation at λ = 458 nm^30^. First, to check the response of microtubules to Rac1 activation, we transfected MDCK cells with mCherry PA-Rac1 and labeled microtubules with SiR-tubulin. On photoactivation of Rac1, bundled microtubules opened up and grew perpendicularly to the cell edge (Fig. 3e, Supplementary video 5), while we did not observe such microtubule dynamics in the neighboring nonactivated cells (Figs. 3e, Supplementary video 5). Subsequently, we co-transfected MDCK cells with mCherry-PA-Rac1 and GFP-Sec61β to study ER dynamics. On photoactivation of Rac1, ER matrices converted to tubules at the site of activation of Rac1, while ER morphology did not change significantly in the non-activated region of the cell (Fig. 3f, Supplementary video 6). Quantifying ER tubule fraction on Rac1 activation showed a significant increase in this parameter in activated regions as opposed to nonactivated regions (Fig. 3g). These results together confirmed that the protrusive forces generated due to actin polymerization and branching at the convex edge guide the organization of ER and microtubule network. Conversely, to augment actomyosin contractility, we went on to treat MDCK mApple-Sec61β cells with Calyculin A. Calyculin A is a myosin light chain phosphatase inhibitor, which leads to hyperactivated myosin^31^. We observed a considerable expansion of ER sheets and matrices in the cell on Calyculin A treatment (Fig. 3h) and a decrease in ER tubule fraction (Fig. 3i).

Next, we explored whether these mechanical forces influence ER structure in the *ex vivo* wound healing model. To test this, we pretreated the embryonic skin explants with Blebbistatin (100 μM) or CK666 (100 μM) two hours before wounding. After creating the wounds, we let the migration happen in the medium containing the inhibitors. We then fixed the tissue and stained it for KDEL and Climp63. We observed that on Blebbistatin-treatment, the cells at both convex and concave edges had considerably less accumulation of Climp63 as compared to control (Fig. 3j) whereas on CK666-treatment, we observed high Climp63 signal at the edge of the cells at both convex and concave edge (Fig. 3j). Taken together, these results suggested that the formation of ER tubules and sheets at convex and concave edges, respectively, depends on the different cellular forces acting at different edge curvatures.

### Mathematical model of edge curvature-dependent ER morphologies

Next, to obtain a fundamental understanding of why the endoplasmic reticulum reorganization takes place in response to geometrical cues, we constructed a mathematical model. We assume that ER behaves like a flexible fiber and its morphology-dependent deformation contributes to the strain energy of the cell. We hypothesize that the difference in the morphologies of ER observed at convex and concave edges are due to differences in the strain-energy state of the cell when the cell contracts and protrudes. Specifically, we tested whether ER tends to be in a tube-like morphology during protrusion and sheet-like during contraction because the combination of morphologies and forces acting on the cell results in lower strain energy.

We modelled cells as 3D mechanical entities with ER embedded in the cytoplasm, and an actin cortex present at the cell edges. The geometry considered for analysis was the front of the cell where ER morphologies were observed to change due to varying mechanical and geometric stimuli. Hence, we did not take the effect of the nucleus into account. The actin cortex was assumed to generate the forces necessary for either protrusion or contraction, depending on the curvature. We assumed cytoplasm as a linear elastic solid while ER fibers were modelled as linear elastic beams. The actin cortex was modelled with a strain-rate dependent non-linear constitutive law, as proposed previously^32^. *In vitro* and *ex vivo* experiments showed that ER exhibited a tube-like morphology at the convex edge during lamellipodia-based protrusion and sheet-like morphology at the concave edge during purse-string movements. In-silico studies are performed by considering cells with convex or concave edges with varying magnitude of curvatures and ER having either rectangular (sheet-like) or circular (tube-like) cross sections with actin cortex generating different cellular forces based on the edge curvature. We compared the strain energy density, as given in

Eq. (*1*), between the cases to find out which morphological state of ER was preferred for the given edge curvature, where σis the stress, ϵ is the strain and V is the volume of the cell.

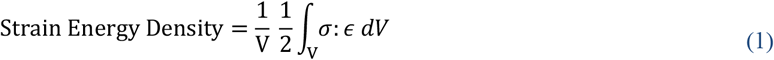

To begin with, we considered a cell with a flat edge denoted as having a curvature of 0 (Fig. 4a, Supplementary Fig. 5a). We observed that when cells experienced protrusive forces, the cell that had ER in the tubular state and aligned along the direction of the protrusion, exhibited lower strain energy density compared to ER aligned perpendicular to protrusion (Supplementary Fig. 5b). ER aligning along the direction of protrusion experienced mainly axial forces with very low bending moments. The high axial stiffness of ER resulted in lower deformation of the cell and thereby lowered the strain energy density. In contrast, during contraction, cells experience higher bending moments compared to axial forces. Hence, a cell with ER aligned parallel to the cell edge exhibited lower strain energy density compared to ER aligned perpendicular to the cell edge (Supplementary Fig. 5b).

**Figure 4.**
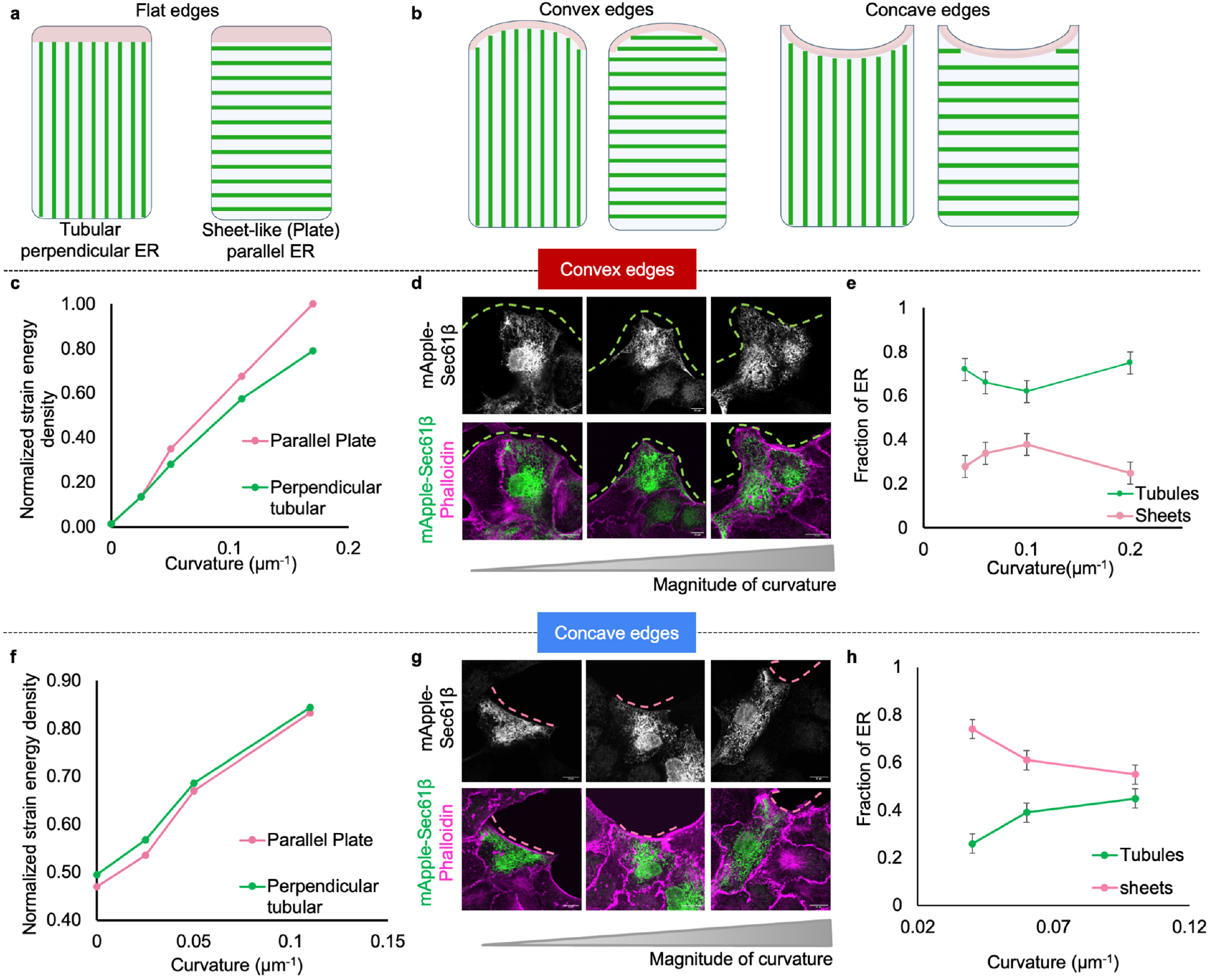
Strain energy minimization promotes different ER morphologies. **a**, Schematic representing front of the cell with straight edge, actin cortex (magenta) and ER (Green) with perpendicular (left) and parallel (right) orientation **b**, schematic representing front of the cell at positive (left) and negative (right) curvature with ER in perpendicular and parallel orientations, **c**, Normalized strain energy density of parallel (pink) and perpendicular (green) ER with increasing positive curvature during protrusion **d**, Representative images of MDCK cells expressing mApple-Sec61β (green) and stained with phalloidin (magenta) migrating at positive curvatures of increasing magnitude (left to right) scale bar-10μm **e**, quantification of fraction of tubules at front of the cells migrating at varying magnitude of positive curvature (n=50) **f**, Normalized strain energy density of parallel (pink) and perpendicular (green) ER with increasing negative curvature during contractility **g**, Representative images of MDCK cells expressing mApple-Sec61β (green) and stained with phalloidin (magenta) migrating at negative curvatures of increasing magnitude (left to right) scale bar-10μm **h**, quantification of fraction of tubules at front of the cells migrating at varying magnitude of concave curvature (n=42). Pink dashed lines mark the negative curvatures, green dashed lines mark the positive curvatures **(d**,**g)**

Next, we added positive (convex) and negative (concave) curvatures to the cell (Fig 4b). For a given curvature, we kept the volume of ER in the cell constant between different ER morphologies considered. We observed that during protrusion at positive curvature, the strain energy density of a cell with parallel ER (sheet-like) was higher than that of a cell with perpendicular ER (tube-like) indicating that ER prefers to stay in the tube-like morphology aligned along the direction of protrusion (Fig. 4c). In addition, as the curvature increased (Supplementary Fig. 5c), the difference in strain energy density between cells with parallel ER and perpendicular ER morphologies increased further (Fig. 4c). Plotting the displacement profile for the cell undergoing protrusion with ER sheets and tubules (Supplementary Fig. 5d) showed that the displacement at the tip was higher for cell with sheet morphology compared to tubular morphology, leading to higher elastic strain energy. Based on this theoretical result, we hypothesized that as the curvature of the cell increased, ER further aligned along the direction of protrusion. To test this hypothesis experimentally, we calculated the amount of ER tubules and sheets at the front of the cells at positive curvatures of different magnitudes. As predicted by the model, we observed that as the curvature increased, the amount of tubules at the front increased and the amount of sheets decreased (Figs. 4d-e).

In contrast, during contraction, strain energy density with parallel ER was lower than that of a cell with perpendicular ER, indicating that cells preferred parallel ER morphology over perpendicular ER morphology (Fig. 4f). But, as the curvature increased, the difference in energy density between parallel and perpendicular morphologies decreased (Fig. 4f). In contrast to protrusion, under contraction, displacement profiles showed that cell with ER tubule morphology had higher displacement compared to cell with sheet morphology (Supplementary Fig. 5e). We hypothesized that with an increase in curvature, due to increase in axial forces, perpendicular ER morphology might be present in cells along with parallel ER morphology. Therefore, we quantified ER tubule and sheet contents at varying magnitudes of negative curvature, and indeed, we observed that at low negative curvature, on average the cells contained 75% ER sheets at the front and as the magnitude increased, the amount of sheets decreased and tubules increased (Figs. 4g-h). Collectively, from the simulation results shown, we inferred that the probability of finding a particular distribution of ER depends on the curvature of the cell, and whether the cell is undergoing contraction or protrusion. Therefore, the distribution of ER responds to the geometric and biomechanical stimuli present in a cell.

### ER-microtubule interaction promotes ER tubule generation at the convex edge

ER and microtubules have significant structural interdependency^27,33-35^. Moreover, the ER morphology and dynamics are known to be regulated by microtubules^35,36^. In our study, we observed a great correlation between the organization of ER and microtubules. The tubular ER network coincided with the perpendicularly arranged microtubules, while lamellar ER coincided with parallelly arranged microtubules (Supplementary Fig. 6a). We therefore wondered whether microtubules might guide edge curvature-dependent ER morphology. To this end, we first disrupted microtubules by treating the cells with 5 μM nocodazole. As expected^37^, upon nocodazole-treatment, ER tubules disappeared irrespective of the edge curvature (Figs. 5a-b). We then tested whether ER tubule formation at the convex edge involved ER movements along microtubules by sliding and tip attachment complex (TAC) dynamics^38,39^ (Fig. 5c). We perturbed the ER sliding dynamics by expressing KHC+ (Kinectin binding domain of Kinesin) or KNT+ (Kinesin binding domain of Kinectin), which led to disruption of the Kinectin-Kinesin interaction^40^. In the cells expressing KHC+ or KNT+, the fraction of ER tubules at the convex edge was significantly reduced as compared to the control EGFP-expressing cells (Figs. 5d-e). But at the concave edge, the accumulation of ER sheets in KHC+ and KNT+ expressing cells did not change as compared the control EGFP-expressing cells (Figs. 5d-f). Next, we perturbed the tip attachment complex movement of the ER by expressing a dominant negative form (EB1c) of microtubule plus end-binding protein1, EB1, which led to its reduced binding with the microtubules^41^. In mutant EB1c-expressing cells, the fraction of ER tubules at the front decreased at the convex edge (Figs. 5g-h) but the accumulation of ER sheets at the concave edge did not display any change as compared to the control cells (Figs. 5g-i). Taken together, these results suggested that both Kinectin-Kinesin dependent sliding and tip attachment complex (TAC) dynamics of ER are important for the generation of ER tubules at the convex edge but not for ER sheet accumulation at the concave edge.

**Figure 5.**
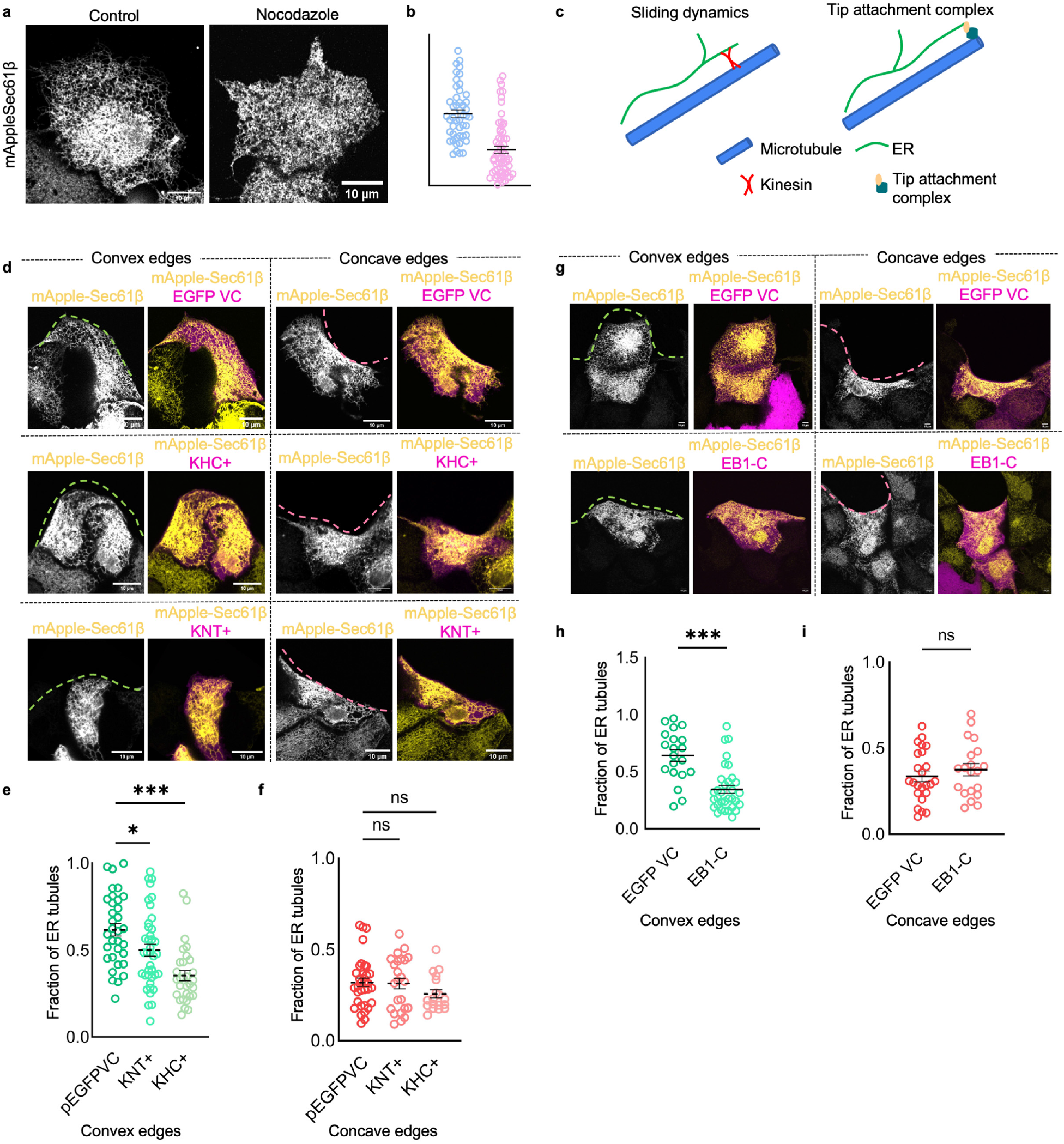
ER-microtubule interaction is important for ER organization at the convex edge. **a**, Representative images of MDCK expressing mApplesec61β treated with DMSO (left) and Nocodazole (right) scale bar -10μm **b**, quantification of fraction of tubules in the whole cell treated with DMSO or nocodazole n=50, 68 from left to right **c**, schematic representing sliding and TAC dependent movement of ER on microtubules **d**, representative images of MDCK cells expressing mApplesec61β (yellow) and pEGFP control vector (magenta, upper panel) or KHC+ (magenta, middle panel) or KNT+ (magenta, lower panel) migrating at convex (left) and concave (right) edge scale bar-10 μm. **e**,**f** quantification of fraction of tubules at the front of cells expressing pEGFP vector control, KHC+ or KNT+ migrating at **e**, convex edge n=33, 39, 30 from left to right and **f**, concave edge n=34, 24, 18 from left to right **g**, Representative images of MDCK cells expressing mApplesec61β (yellow) and EGFP vector control (magenta, upper panel) or EB1-c (magenta, lower panel) migrating at convex (left) and concave (right) edge, scale bar-10μm **h**,**i** quantification of fraction of tubules at the front of the cells expressing EGFP vector control or EB1-c migrating at **h**, convex edge n= 20, 34 from left to right **i**, concave edge n=24, 20 from left to right. Data are mean ± s.e.m. Anova tests **(e**,**f)**, two tailed t-test **(h**,**i).** Pink dashed lines mark the concave edges, green dashed lines mark the convex edges **(d**,**g)**

### Manipulating ER morphology alters the cellular response to edge curvature

Next, we determined whether different ER morphologies at different edge curvature might be consequential for cells at the edge determining the curvature-specific mode of epithelial migration. To this end, we manipulated the ER morphology by overexpressing either an integral ER membrane protein Rtn4a, which leads to the enrichment of ER tubules, or an ER sheet-associated protein Climp63, which leads to the enrichment of ER sheet^42^. Indeed, MDCK cells overexpressing Rtn4a-GFP showed higher ER tubule fraction and MDCK cells overexpressing mCherry-Climp63 showed higher ER sheet fraction as compared to the control cells overexpressing mApple-Sec61β (Supplementary Fig. 7a). Subsequently, we seeded these cells around the PDMS micropattern and studied their migration behavior at different edge curvatures. To normalize across experiments, we calculated the probability of forming lamellipodia by dividing the number of transfected cells forming the lamellipodia by the total number of the transfected cells at a particular edge curvature. For cells overexpressing Rtn4a, the percentage of cells forming lamellipodia at the convex edge was comparable to that for wildtype cells. But at the concave edge, where the wildtype cells majorly formed actin bundles, Rtn4a-overexpressing cells showed a significantly higher probability of lamellipodium formation (Figs. 6a-b). In contrast, Climp63 overexpression, which amplified ER sheet morphology, resulted in a significant decrease in the percentage of the lamellipodium forming cells at the convex edge (Figs. 6c-d). To further confirm the identity of the protrusive structures at the gap edge, we stained the cells with an anti-Cortactin antibody. Cortactin is a molecular marker for lamellipodium and was expectedly present at the protrusion in wildtype cells at the convex edge but not at the concave edge (Supplementary Figs. 7b-c). Interestingly, Rtn4a-overexpression induced Cortactin expression even at the concave edge (Supplementary Fig. 7c) while Climp63-overexpression abolished it at the convex edge (Supplementary Figs. 7b).

**Figure 6.**
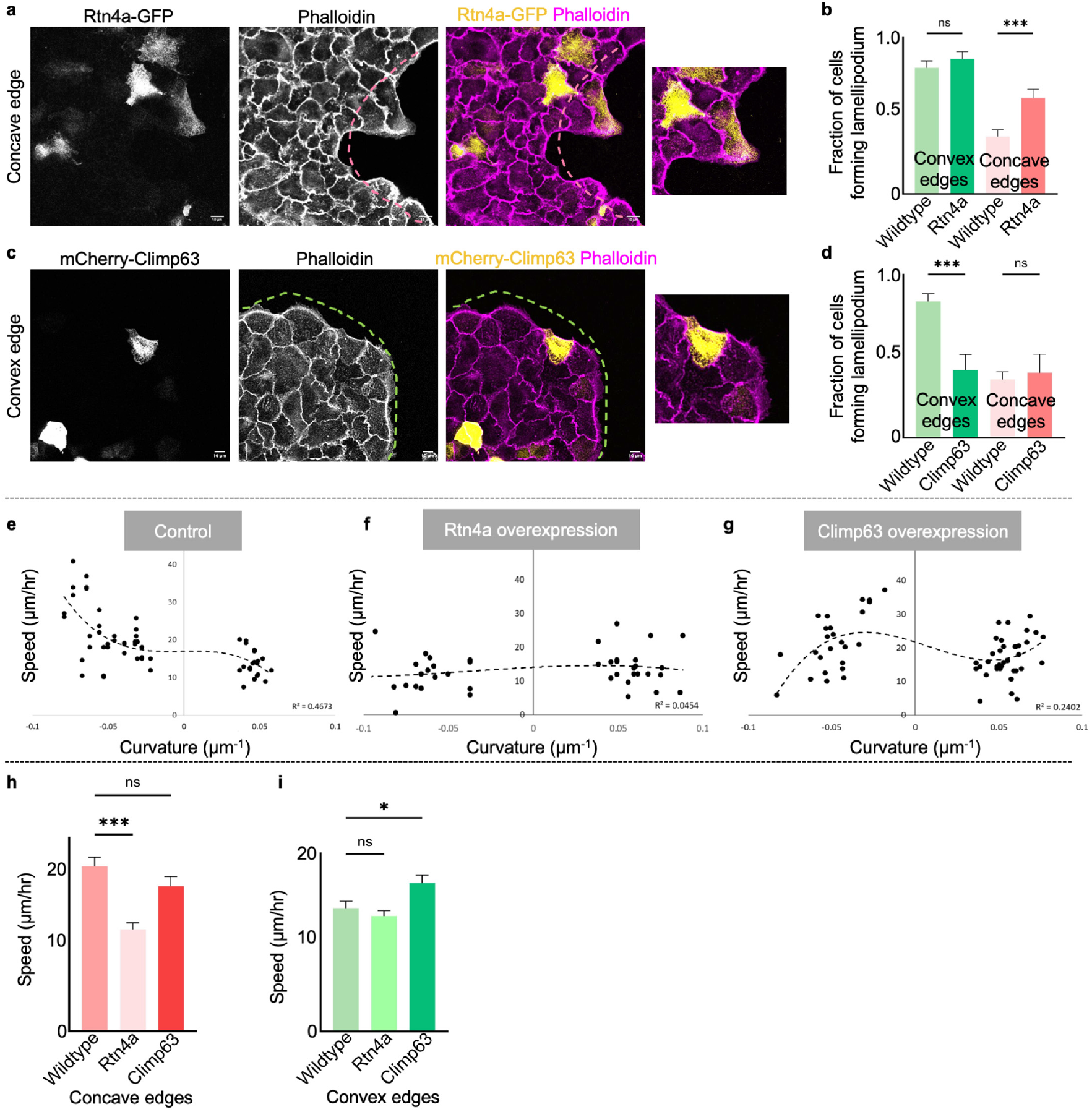
Curvature-specific ER morphologies regulate the modes of epithelial migration. **a**, Representative MDCK cells migrating at concave edge heterogeneously expressing Rtn4a-GFP (yellow), stained with phalloidin (magenta), zoomed inset shows Rtn4a-GFP expressing cell forming lamellipodia at concave edge scale bar-10μm **b**, quantification of fraction of MDCK WT and Rtn4a over-expressing cells forming lamellipodia at convex and concave edges n=100 cells **c**, Representative MDCK cells migrating at convex edge heterogeneously expressing mCherry-Climp63 (yellow), stained with phalloidin (magenta) zoomed inset shows mCherry-Climp63 expressing cell forming actin bundle at convex edge scale bar-10μm **d**, quantification of fraction of MDCK WT and Climp63 over-expressing cells forming lamellipodia at convex and concave edges n=85 cells **e**, Plot of initial velocity of MDCK WT cells as a function of curvature **f**, Plot of initial velocity of MDCK cells expressing Rtn4a-GFP as a function of curvature **g**, Plot of initial velocity of MDCK cells expressing mCherry-Climp63 as a function of curvature **h**,**i** quantification of average velocity of WT, Rtn4a-GFP and mCherry-Climp63 expressing cells at **h**, concave n=39, 42, 44 from left to right and **i**, convex edge n=30, 45, 45 from left to right. Data are mean ± s.e.m. Anova test **(b**,**d**,**h**,**i)** Pink dashed lines mark the concave edges, green dashed lines mark the convex edges **(a**,**c)**

Previous studies have shown that the different migration modes at different edge curvatures also affect the speed of collective cell migration, with cells migrating less rapidly with increasing edge curvature^15^. We calculated the initial speeds of control, Rtn4a-overexpressing, and Climp63-overexpressing cells at different edge curvatures. As expected, wildtype cells moved faster at the concave edge than at the convex edge (Fig. 6e). However, in cells overexpressing Rtn4a, the edge curvature-dependent difference in speed was lost, and cells migrated at similar speeds at both edge curvatures (Fig. 6f). Similarly, in Climp63-overexpressing cells, the initial speed did not change as a function of the edge geometry (Fig. 6g). Next, we computed the average initial velocity of wildtype, Rtn4a-overexpressing, and Climp63-overexpressing cells at different edge curvatures. At the concave edge, Rtn4a-overexpressing cells migrated slowly as compared to the wildtype cells (Fig. 6h). At the convex edge, Climp63-overexpressing cells migrated at a higher pace as compared to the wildtype cells and Rtn4a cells (Fig 6i). Together, these results suggested that epithelial cells at the edge may choose a curvature-specific mode of migration based on the varying morphologies of the ER at different edge curvatures.

### ER morphologies regulate edge curvature-specific orientations of focal adhesions

Since ER-resident proteins interact with the focal adhesion complex^43-46^, ER dynamics might be crucial for the maturation of cell-substrate adhesions during cell adhesion and migration^47^. Relevantly, focal adhesions orient differently at the two edge curvatures during migration, aligning perpendicular to the edge at convex regions and parallel to the edge at concave regions^15,16^. We, therefore, hypothesized that different ER morphologies at the convex and concave edges might influence the orientation and stability of focal adhesions. To test this hypothesis, we first checked whether ER tubules or sheets contacted focal adhesion punctae, using Paxillin as the focal adhesion marker. Co-localization images revealed that ER tubules were associated with perpendicular Paxillin punctae at the convex edge, while ER sheets were associated with parallel focal adhesions at the concave edge (Fig. 7a). Next, we quantified the fraction of focal adhesions of a particular orientation associated with ER sheets or tubules. Adhesions aligned at 0-18 degrees relative to the edge were considered as parallel and those aligned at 72-90 degrees were considered as perpendicular focal adhesions. Quantification revealed that at both curvatures, perpendicular focal adhesions mainly contacted ER tubules, while the parallel focal adhesions were associated with the ER sheets (Fig. 7b). Then, to determine whether altering ER morphology would alter the orientation of focal adhesions, we examined the orientation of Paxillin punctae in Rtn4a- or Climp63-overexpressing cells (Fig. 7c), which showed an increased propensity to form ER tubules and sheets, respectively. Subsequently, Rtn4a-overexpression led to a decrease in the fraction of parallelly oriented focal adhesions and increased perpendicularly oriented adhesions at the concave edge (Fig. 7c). In contrast, Climp63-overexpression led to a decrease in the fraction of perpendicularly oriented focal adhesions and increased parallelly oriented focal adhesions at the convex edge (Fig. 7c). Taken together, these results suggested that morphological changes in the endoplasmic reticulum modulate focal adhesion, promoting different modes of migration at the two curvatures.

**Figure 7.**
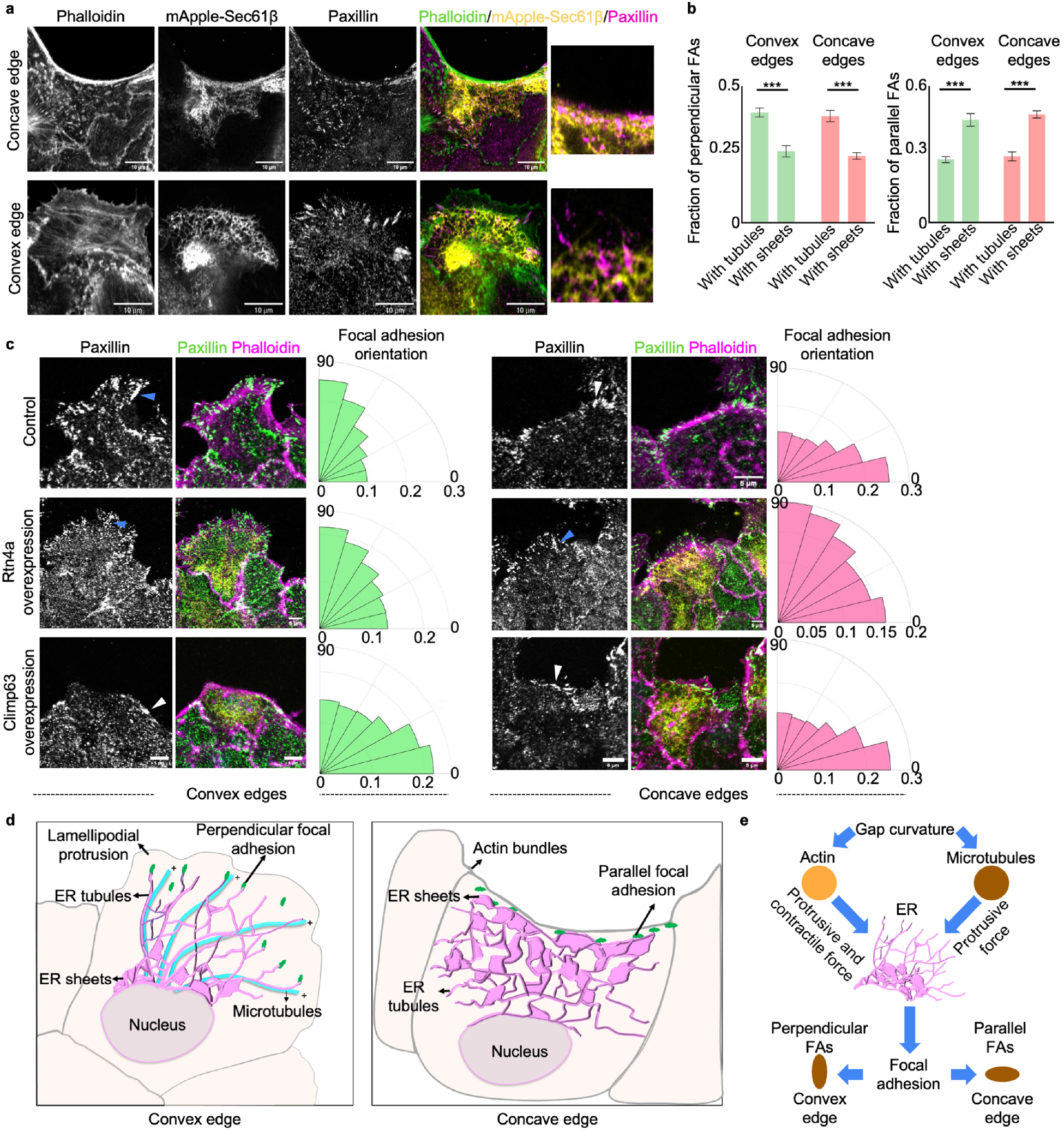
ER morphologies affect focal adhesion orientations. **a**, Representative images of MDCK cells expressing mApple-Sec61β (yellow) migrating at convex (upper panel) and concave (lower panel) edge stained with anti-paxillin (magenta) and phalloidin (green), scale bar -10μm, zoomed insets. Show association between ER tubules and perpendicular adhesions (upper panel) and ER sheets and parallel adhesions (lower panel) **b**, quantification of fraction of perpendicular focal adhesions n= 58, 26, 47, 70 from left to right and parallel focal adhesions n=58, 26, 47, 70 from left to right **c**, Representative images of MDCK WT (upper panel), Rtn4a-GFP (yellow, middle panel), mCherryClimp63 (yellow, lower panel) cells stained with anti-paxillin (green) and phalloidin (magenta), blue arrowhead show perpendicular FA, white arrowhead show parallel FA, Rose plot at the right show quantification for focal adhesion orientation w.r.t wound from 0 to 90 degrees (at convex WT n=63, Rtn4a n= 77, Climp63 n= 103; at concave WT n= 51, Rtn4a n= 100, Climp63 n= 62). **d**, Summary of the study: schematic (left) depicting ER tubules at the front of cells migrating at convex edge sliding and growing along the microtubules, and associating with the perpendicularly oriented focal adhesions promoting lamellipodial crawling. Schematic (right) depicting ER sheets accumulated at the front of cells migrating at the concave edge, associating with parallel focal adhesions and promoting actin bundle-driven purse string closure. **e**, ER in mechanotransduction. Positioning ER as a key player in mechanotransduction, integrating signals from microtubules and actomyosin networks to regulate cellular responses to physical stimuli. Data are mean ± s.e.m. Anova test **(b)**.

## DISCUSSION

Epithelial gap closure is an essential and universal component in physiological and pathological processes and happens throughout the lifespan of an organism. It involves collective cell movement through lamellipodia-mediated crawling and purse-string-like movements, each exhibiting distinct actin dynamics and focal adhesion assembly. The cellular response to gaps was found to strongly depend on edge curvature. However, the mechanisms driving this response remain largely unknown. In this study, we explored how collectively migrating epithelial cells alter their intracellular network of organelles in response to the shape of the gap curvature. Our findings reveal that among all organelles, the endoplasmic reticulum (ER) undergoes conspicuous morphological changes in response to edge curvature and plays a pivotal role in determining the mode of epithelial migration across varying edge curvatures (Fig. 7d). ER is the largest and the most dynamic cell organelle, which has been hypothesized to be a sensor of multiple extracellular cues attributing to its dynamicity and elaborate structure spanning the whole cell^17,24,48,49^. ER exhibiting tubular morphology at the convex edge and sheet accumulation at the concave edge raises the question if the ER organization is involved in sensing and responding to physical and geometric cues, and our work presented strong evidence that this is the case. This gap curvature-dependent behavior of ER appeared consistent across multiple epithelial cell lines with controlled predisposed geometrical cues, as well as in mouse embryonic epidermis, where spontaneous geometric cues arise upon injury. Our experiments further suggest that these changes in ER morphology depend on the protrusive forces, supported by both actin polymerization and microtubules, and contractile forces, supported by actomyosin networks, placing ER at a cross point between two important cytoskeletal elements, namely actin filaments, and microtubules. In the future, it will be interesting to study whether intermediate filaments such as cytokeratin and vimentin play any role in the gap curvature-dependent differential organization of ER. While actin and microtubules play an important role in determining the ER morphologies at different gap curvatures, we also provide a complimentary explanation for specific ER morphologies. A theoretical model trying to understand the ER morphologies from a physical perspective suggests that under protrusive forces at the convex edge, perpendicular ER tubules have lower strain energy whereas under contractile forces at the concave edge parallel sheet ER has lower strain energy. Strain energy minimization, therefore, might underlie different ER morphologies, making ER tubules favored at the convex edge and ER sheets at the concave edge. Hence, future studies should attempt to disentangle the interaction between active and passive mechanisms driving specific ER morphologies.

The interrelationship between ER and microtubules has been explored in many contexts in mammalian cells, and it can be fairly concluded that they are interdependent structures. We show that the formation of ER tubules at the convex edge requires microtubule-dependent ER dynamics via both kinesin-dependent sliding movement and tip attachment complex-mediated movement. However, at the concave edge, these ER-microtubule interactions did not seem to contribute to ER sheet accumulation at the edge. It is still possible that some of the ER sheet-associated proteins such as Climp63 and p180, which possess microtubule binding domains^50,51^, may contribute to the accumulation of ER sheets at the concave edge. Additionally, whether actomyosin contractility can directly promote ER sheet accumulation independent of microtubules remains speculative. On the other hand, ER structure and dynamics have also been shown to influence the distribution and post-translational modifications of microtubules^27,52^. It is tempting to speculate that the reorganization of ER in response to curvature could affect post-transitional modifications of microtubules thereby affecting the overall distribution and polarity of other organelles eventually as the migration progresses.

A functional consequence of ER morphology in the modes of epithelial migration is that the manipulation of ER morphologies at different gap curvatures led to a switch in the mode of migration. Specifically, Climp63-overexpressing cells formed prominent actin bundles even at the convex edge, while Rtn4a-overexpressing cells formed prominent protrusions even at the concave edge. The mechanism by which ER morphology affects cell migration involves focal adhesion orientation. However, there may be additional mechanisms. Here it should be appreciated that ER-mediated regulation of focal adhesions could be just one of the mechanisms by which ER dictates the mode of epithelial cell migration. ER tubules are known to be involved in lipid biogenesis, calcium signaling, and making membrane contact sites with other organelles and the plasma membrane. A reticulated network of ER tubules at the convex edge could also regulate lamellipodial crawling by providing membrane lipids via the contact sites and by releasing calcium for the activation of the actomyosin network. ER sheets on the other hand are studded with ribosomes and are involved in protein synthesis. Whether sheet accumulation at the concave edge leads to directed protein synthesis and delivery facilitating purse-string-like movements remains elusive. How the conventional roles of ER subdomains differ at the two curvatures and how they eventually guide cell migration remains an open question. Despite these unknowns, the intriguing results of our study promote ER as a central coordinator for collective cell migration and open a new frontier for organelle research in cell biology.

On one hand, our experiments reveal that changes in ER morphology in response to gap curvature are influenced by both protrusive forces and contractile forces. On the other hand, ER morphologies play important roles in supporting the curvature-specific mode of cell migration. Overall, our research places the ER within the broader network of mechanotransduction (Fig. 7e), emphasizing its role as a sensor and transducer of mechanical cues. By integrating signals from the microtubules and actomyosin networks, the ER appears to play a central role in regulating cellular responses to physical stimuli, ultimately influencing cell migration processes. This understanding of the intricate interplay between ER morphology and mechanotransduction pathways provides valuable insights into the fundamental mechanisms underlying cell behavior and will have implications for various physiological and pathological processes.

## Notes

### Competing Interest Statement

The authors have declared no competing interest.

### Funding

- Science and Engineering Research Board (SERB) India (now ANRF): CRG/2021/003907

