## Supplementary Material for "Curvature-dependent morphological reorganization of the endoplasmic reticulum determines the mode of epithelial migration"

**Affiliations:**
1. Tata Institute of Fundamental Research Hyderabad (TIFRH), Hyderabad, Telangana 500046, India
2. School of Mathematics, University of Birmingham, Edgbaston, Birmingham B15 2TT, UK

**This document contains:**

A. Materials and Methods
B. Supplementary Table 1: Antibodies and fluorophores
C. Supplementary Table 2: Plasmids
D. Theoretical Modelling
E. Supplementary References
F. Supplementary Figure Legends
G. Supplementary Video Legends
H. Supplementary Figures

### A. MATERIALS AND METHODS

**Cell culture.** The Tetracycline-resistant wild-type Madin-Darby canine kidney (MDCK-WT) cells utilized in this investigation were generously provided by Yasuyuki Fujita. These cells were cultured in Dulbecco's modified Eagle's medium (DMEM) with GlutaMAX (Gibco), supplemented with 5% fetal bovine serum (Tet-free FBS, Takara Bio), and 10 U/mL penicillin and 10 μg/mL streptomycin (Pen-Strep, Invitrogen). Cultures were maintained in a $CO_2$ incubator at 37°C. EPH4-Ev epithelial cells (ATCC; CRL-3063), employed in the study, were maintained in DMEM with GlutaMax (Gibco), supplemented with 10% fetal bovine serum (Tet-free FBS, Takara Bio), and 1.2 μg/mL puromycin (Gibco). These cells were also maintained in a 37°C, 5% $CO_2$ environment. Subculturing of both cell lines occurred every 3 to 4 days utilizing Trypsin/EDTA (Invitrogen). For transfection, cells were subjected to Lipofectamine 2000 (Invitrogen) following the manufacturer's instructions. Approximately 8 to 10 hours post-transfection, cells were seeded for the migration assays.

**Micropatterning.** Polydimethylsiloxane (PDMS) stencils with pillars of defined shapes were created using soft photolithography. Briefly, 150 μm thick SU-8 2075 (Kayaku advanced materials, Y111074) was coated on a silicon wafer followed by soft baking at 65°C for 5 minutes and 95°C for 20 minutes. SU-8 was exposed in the desired pattern using the MicroWriter ML 3 Pro (Durham Optics Magneto Ltd) followed by post-exposure baking at 65°C for 7 minutes and 95°C for 15 minutes. The unexposed photoresist was removed by immersing the wafers in the SU-8 developer (Kayaku advanced materials, Y020100) for 8-10 minutes. Patterned SU-8 wafers were then hard-baked at 170°C for 45 minutes. PDMS (Sylgard 184, Dow Corning) was mixed in a ratio of 1:10, degassed, and poured over the patterned wafers. After curing at 85°C for 2 hours, stencils with pillars were peeled off. The stencils were then treated with 2% Pluronic acid for 2 hours followed by 15 minutes of 1X PBS wash. The PDMS stencil and a petri dish were plasma-cleaned for 10 seconds, and the stencil was then placed on the dish. Cells were seeded around the pillars, and after confluence, the stencils were removed allowing the cells to migrate into the gaps.

**Cell migration assay.** A solution of 20 μg/mL fibronectin was applied to a glass-bottomed petri dish (IBIDI). After three washes to remove excess protein, the dish was air-dried. The PDMS stencil underwent plasma cleaning for 20 seconds before being positioned onto the dish. The petri dish with the stencil was then subjected to UV sterilization for 1 hour. A concentrated suspension of MDCK or other cells was seeded around the pillars, followed by the addition of excess media around the stencil. Cells were allowed to settle around the pillars for 15-18 hours. Subsequently, the stencils were lifted, enabling cell migration at various curvatures. Live cell imaging or fixation occurred either immediately or after 30 minutes of migration.

**Mouse embryonic skin wounding.** E14.5 - E16.5 mouse embryos were dissected out. The dorsal skin from these embryos was then excised. The skin was placed on a filter paper (Whatman filter, 8 μm pore size) with the epidermal side up. Using a scalpel/fine scissor, incision wounds were made on the epidermis. The filter paper along with the tissue was kept floating in a 6-well plate containing 2 mL of DMEM with 5% fetal bovine serum (FBS) and 1% Pen-Strep. After the desired incubation period, the tissue was fixed in 4% paraformaldehyde (PFA) for 2 hours at room temp. The tissue was washed in PBS and transferred to 90% ethanol in PBST (0.2% Triton X-100 in 1X PBS) overnight at -20°C. The tissue was then rehydrated using graded ethanol (75%, 50%, and 25% in PBST). After two more washes in PBST, the tissue was blocked using 10% normal goat serum (NGS) in PBST for 2 hours. Primary antibodies were added and incubated overnight at 4°C. This was followed by subsequent PBST washes, after which secondary antibodies were added and incubated for

2 hours at room temperature. After washing, the tissue was mounted on a slide using Fluoroshield (Sigma Aldrich) mounting media. The slide was cured overnight and imaged using an upright confocal (Leica Stellaris). For inhibitor experiments, the tissue was incubated with the inhibitor, Blebbistatin (100 μM) or CK666 (100 μM) in the complete medium for two hours before making the incision wounds. After 2 hours, the wounds were made, and the tissue was incubated in the same media for the next 30 minutes followed by fixation and antibody staining.

**Immunofluorescence.** Cells were fixed using a solution containing 3% formaldehyde and 0.1% glutaraldehyde diluted in 1X PBS for 10 minutes at room temperature. Following fixation, cells underwent three washes with 1X PBS. Subsequently, permeabilization was carried out using 0.25% Triton X-100 in PBS for 10 minutes at room temperature, followed by three quick washes with 1X PBS. Cells were then incubated with a blocking solution (consisting of 0.1% Triton X-100 in 1X PBS + 2% BSA) for 1 hour at room temperature. Afterward, cells were incubated with a primary antibody prepared in the blocking solution for 2 hours at room temperature. Following the primary antibody incubation, cells were washed thrice with 1X PBS and then incubated with the secondary antibody, DAPI (1:1000), and Phalloidin (1:40) dilutions, all prepared in the blocking/staining solution, for 1 hour at room temperature. Finally, the samples underwent two quick washes and a 5-minute wash with 1X PBS before proceeding to imaging.

**Confocal microscopy.** Fluorescence images were captured using a 60X oil objective (PlanApo N 60x Oil, NA=1.42, Olympus) and a 100X oil objective (UPlanSApo, 100X/1.40 oil), both mounted on an Olympus IX83 inverted microscope equipped with a scanning laser confocal head (Olympus FV3000). Time-lapse imaging of live samples was conducted in the live-cell chamber provided with the microscopy setup. For the photoactivation experiment, a 60X oil objective (PlanApo N 60x Oil, NA=1.42, Olympus) mounted on an Olympus IX83 inverted microscope equipped with a scanning laser confocal head (Olympus FV3000) was utilized, supported by a live-cell imaging setup. MDCK cells, either co-expressing Photoactivable Rac1 and GFP-Sec61β or expressing photoactivable Rac1 and labeled with SiR-tubulin, were seeded on a glass-bottom dish in OptiMEM medium with HEPES (20 mM). A region of interest approximately 5 μm in diameter was irradiated with a 1% intensity 445 nm laser at 1000 μs/pixel, followed by LSM imaging of the two channels (561 nm and 488 nm or 561 nm and 647 nm) at 0.1-0.5% intensity. This cycle of activation and imaging was repeated continuously for 15-20 minutes.

**SoRa (Super-resolution by optical Reassignment).** Fluorescence images were acquired using 100X oil objective (Apo TIRF, NA=1.49, WD=0.12) mounted on a Nikon inverted research microscope Eclipse T*i2*-E with Yokogawa CSU-W1 SoRa unit. A further 2.8X magnification was employed to acquire super-resolution images with the SoRa unit. After the acquisition, images were subjected to Richardson-Lucy 3D deconvolution.

**Inhibition studies.** For all inhibition studies, cells underwent pretreatment with the desired inhibitor concentration in complete DMEM for 1 hour at 37°C in a 5% $CO_2$ humidified incubator before lifting the PDMS stamps. Throughout the migration process, cells remained in complete media containing the inhibitor at the specified concentration for the required migration duration. To modulate actomyosin contractility, MDCK cells were treated with Blebbistatin (Sigma), a myosin inhibitor, at a concentration of 50 μM. Arp2/3-inhibition was achieved using CK666 (Sigma) at a concentration of 50 μM. Calyculin A (Sigma), a phosphatase inhibitor, was employed to enhance actomyosin contractility at a concentration of

20 nM. Nocodazole (Sigma) was utilized to disrupt microtubules in MDCK cells at a concentration of 10 μM. Following inhibitor treatment, cells were fixed and immunostained with the desired antibodies.

**Antibodies and plasmids.** Source and dilution information for all primary and secondary antibodies used in immunofluorescence staining are given in Supplementary Table 1. Details of plasmids used in this study are listed in Supplementary Table 2 with their source.

**Image analysis.** Image analysis was conducted using Fiji. To determine the fraction of ER tubules, trainable Weka segmentation was utilized to distinguish between tubules and sheets within the ER structures. The ER tubule fraction at the leading edge of the cell was computed by dividing the tubular ER area by the total ER area manually marked in the region at the migrating front of the cell. The same classifier was applied for ER segmentation in all inhibition and mutant studies. In the photo-activation experiment of Rac1, the region of interest (ROI) for Rac1 activation was selected for segmentation, while a similarly sized ROI at the non-activated region within the cell served as a control. Microtubule orientation was determined using the directionality plugin in Fiji, with 10 bins and angles ranging from 0 to 180 degrees. An ROI at the front of the migrating cell in the α-tubulin channel was chosen, and directionality frequency was calculated. Similarly, focal adhesion (FA) orientation was analyzed using the directionality plugin with 5 bins and angles ranging from 0 to 90 degrees. An ROI at the cell's leading edge in the paxillin channel was selected, and directionality frequency was computed. FAs aligned between 0-18 degrees were considered parallel to the wound, while those aligned between 72-90 degrees were deemed perpendicular. FAs falling within these ranges were excluded from the analysis. Curvature analysis was performed using the Fiji plugin kappa-curvature analysis.

**Statistical analysis.** Statistical analyses were performed using GraphPad Prism 9. Statistical significance was determined using an Unpaired t-test with Welch's correction or One-way ANOVA, as specified in the corresponding figure legend. Scatter-bar plots were presented as mean ± standard error of the mean (s.e.m.). P-values above 0.05 were considered not statistically significant. Quantification was based on data from a minimum of three independent biological replicates.

### B. Supplementary Table 1: Antibodies and Fluorophores

| S.no. | Antibody/Fluorophore | Catalogue No. | Dilution/Working conc. |
|---|---|---|---|
| 1. | Alexa Fluor 647 Phalloidin | 8940 (CST) | 1:40 |
| 2. | Alexa Fluor 488 Phalloidin | 8878S (CST) | 1:40 |
| 3. | DAPI (4’,6-diamidino-2-phenylindole) | D1306 (Invitrogen) | 1 µg/ml |
| 4. | Anti-α-tubulin | 3873S (CST) | 1:200 |
| 5. | Anti-LAMP1 | Ab24170 (Abcam) | 1:500 |
| 6. | Anti-GRASP65 | MA5-25148 (Thermo Fischer Scientific) | 1:200 |
| 7. | Anti-CKAP4 | 16686-1-AP (Proteintech) | 1:200 |
| 8. | Anti-KDEL | 420400 (Merck) | 1:200 |
| 9. | Anti-Cortactin | SAB5701694 (Merck) | 1:200 |
| 10. | Anti-Paxillin | Ab32084 | 1:200 |
| 11. | Anti-Sec61β | PA3-015 (Thermo Fischer Scientific) | 1:200 |
| 12. | Draq5 | 4084S (CST) | 1:1000 |
| 13. | Mito Tracker Green | 9074S (CST) | 200 nM |
| 14. | Anti-rabbit IgG, AlexaFluor 568 | A11036 (Invitrogen) | 1:500 |
| 15. | Anti-rabbit IgG, AlexaFluor 488 | A11008 (Invitrogen) | 1:500 |
| 16. | Anti-mouse IgG, AlexaFluor 488 | A11001 (Invitrogen) | 1:500 |
| 17. | Anti-mouse IgG, AlexaFluor 568 | A11031 (Invitrogen) | 1:500 |
| 18. | SiR-tubulin | CY-SC006 (Cytoskeleton) | 200 nM |

### C. Supplementary Table 2: Plasmids

| S.No. | Plasmid | Source |
|---|---|---|
| 1. | mApple Sec61β-C1 | 90993 (Addgene) |
| 2. | paRac1 | 22027 (Addgene) |
| 3. | GFP Sec61β | 121159 (Addgene) |
| 4. | pEGFP-KHC+ | Gift from Hanry Yu |
| 5. | pEGFP-KNT+ | Gift from Hanry Yu |
| 6. | pEGFP-Vector | Gift from Hanry Yu |
| 7. | EGFP-EB1c | Gift from Anna Akhmanova |
| 8. | EGFP-Vector | Gift from Anna Akhmanova |
| 9. | Rtn4a-GFP | 61807 (Addgene) |
| 10. | mCherry-Climp63 | 136293 (Addgene) |
| 11. | pYFP-Paxillin | 50543 (Addgene) |

**D. Theoretical modelling**

The endoplasmic reticulum (ER) network is a complex dynamic structure within the cell. Mathematical models have been developed in the past to understand the shape of the ER network using the principles of Langevin dynamics[1, 2]. Further, to study the abundance of tubular and sheet ER for a given membrane curvature, models were developed that study the role of curvature-stabilizing proteins in minimizing the energy of the system[3, 4]. However, our experiments showed the role of protrusive and contractile forces in the regulation of ER morphology. A theoretical understanding of the role of mechanical stimuli on ER morphology is still incomplete. Hence, we developed a computational model of the cell made up of 3 components, cytoplasm, actin cortex and ER. We calculated the strain energy of the cell to determine the preferred ER morphology that is associated with a minimum of the strain energy. Since the ER morphological changes were observed near the cell edges, we neglect the nucleus, and consider the actin cortex to exert protrusive and contractile forces on the ER. Since the purpose of the model is to investigate the effect of different organizations of the ER, we assumed, for simplicity, that the cytoplasm and ER are linear elastic material with the ER modelled as beams embedded in the cytoplasm. The actin cortex is modelled to apply either contractile or protrusive forces depending on the edge curvature through a non-linear constitutive law. The stress in the actin cortex is split into active and passive components. Active stress ($\sigma^a$) is assumed to be strain-rate dependent, and passive stress ($\sigma^p$) is assumed to be linear elastic for simplicity. The total stress tensor ($\boldsymbol{\Sigma}$) in the actin cortex can therefore be written as the sum of active ($\boldsymbol{\sigma}^a$) and passive ($\boldsymbol{\sigma}^p$) stress tensors, as given in Eq. (1)

$$\boldsymbol{\Sigma} = \boldsymbol{\sigma}^a + \boldsymbol{\sigma}^p \qquad (1)$$

The actin cortex is assumed to be made up of actin fibers uniformly distributed over the domain. The active stress in each of these fibers is assumed to depend on its strain rate ( $\dot{\epsilon}\ (\phi, \omega)$) as given in Eq. (2).

$$\sigma^a(\phi, \omega) = \eta(\phi, \omega)\sigma_{max}(1 + \frac{k_v\ \dot{\epsilon}(\phi,\omega)}{\sqrt{1+k_v\dot{\epsilon}(\phi,\omega)^2}}) \qquad (2)$$

where $\phi, \omega$ are the standard angular representation in the spherical coordinate system and $\eta(\phi, \omega)$ indicates the actin fibre concentration at an angle $\phi, \omega$. $\sigma_{max}$ is the maximum stress that the actin fiber can handle, $k_v$ is a constant. Furthermore, the growth of fibre concentration follows an ODE as given in Eq. (3)

$$\dot{\eta}(\phi, \omega) = (1 - \eta(\phi, \omega))Ck_f - \left(1 - \frac{\sigma^a(\phi, \omega)}{\eta(\phi, \omega)\sigma_{max}}\right)\eta(\phi, \omega)k_b \qquad (3)$$

$C$ indicates the calcium concentration, which initiates the fiber formation, while $k_f$ and $k_b$ represent the rate of association and dissociation of fibres respectively. The active stress evaluated in each of the fibres is then homogenized following our previous study[5] to evaluate the active stress tensor. The system of equations is solved using the commercial finite element solver Abaqus 2021 [Dassault systèmes, Simulia Corp]. Solving the mechanical equilibrium results in the contractility of a cell. Hence to generate protrusion, we change the sign of active stress, while, for simplicity and comparability, we fix the magnitude. Thus, depending on the edge curvature, and whether the cell is either under protrusion or contraction, the direction and magnitude of the force acting on the ER is affected. Accordingly, we perform simulations by changing the geometry of the cell, and ER morphologies and compare the strain energy density between the cases to find out which morphological state of ER is preferred for a given condition. Furthermore, we found that during contraction, the strain energy density of a cell with ER having a circular cross-section was slightly higher than that of rectangular cross-section. Hence, in all the simulations, we use ER sheets with rectangular cross section and ER tubules with circular cross section. To be able to have a fair comparison, we divide the strain energy by the volume of the cell, referred to as the strain energy density, evaluated as

$$Strain\ Energy\ Density = \frac{1}{V}\frac{1}{2}\int_V \sigma{:}\,\epsilon\ dV \qquad (4)$$

During the comparison of ER morphologies, we keep the volume of ER the same between the cases to remove any bias that the amount of ER can introduce on the overall stiffness of the cell. Following previous studies[6,7], the Youngs modulus of individual ER fiber is taken as 19.3MPa with a Poisson ratio of 0.45, while that of epithelial cells is taken as 73 kPa with a Poisson ratio of 0.45 [8]. The parameters needed for the growth of active stress are taken from a previous study[9]. Representative figures showing the deformation of the cell during protrusion and contraction are given in Supplementary Fig. 5. The curvature presented in the computational model is the curvature of the ellipse at the inflexion point of the edge evaluated following Eq. (5)

$$\kappa = \frac{ab}{\left(\sqrt{a^2\, sin^2\, t + b^2\, cos^2\, t}\,\right)^3} \qquad (5)$$

where a and b are the axes length of the ellipse and t is the angle at which the curvature is evaluated. The code is made available as open source, available to download from the GitHub page : https://github.com/bkprdp/Curvature_Dependent_ER_Morphology.

**F. Supplementary figure legends**

**Supplementary figure 1. Organelle morphology and polarity in response to wound geometry in MDCK. a,** Quantification of the fraction of cells forming lamellipodia out of the total cells at the edge at two curvatures **b,** color wheel representation of the directionality of microtubules at convex (left) and concave (right) curvatures, color wheel ranging from 0 to 180 represents the angle of microtubules relative to the edge of the wound **c,** quantification for directionality of microtubules at convex (green) and concave (pink) curvature (n= 86 at each curve) **d,** representative images of MDCK cells stained for Golgi, lysosome, and mitochondria at convex (upper panel) and concave (lower panel) curvature. MDCK cells stained for left panel- anti-GRASP65 ( magenta, Golgi marker), phalloidin (green) and DAPI (blue); middle panel- anti-LAMP1 (magenta, lysosome marker), phalloidin (green) and DAPI (blue); right panel- cells labeled with Mitotracker green (magenta, mitochondria marker) and DRAQ5 (blue) **e,** Schematic representing MDE (mean distribution from the edge), the yellow dotted line represents the distance between the center of mass of the cellular entity and the edge of the cell; this distance normalized by the radius of the cell and is referred as MDE **f,** Quantification of MDE for actin, microtubules, ER, Nucleus, Golgi, lysosomes, and mitochondria at convex (green) and concave (pink) curvatures. n = 74, 62, 62 69, 83, 49, 81, 71, 63, 61, 80, 78, 85, 88 from left to right, Data are mean ± s.e.m. ANOVA test.

**Supplementary figure 2. Organelle organization and polarity in response to wound geometry in Eph4 cells. a,b,c** Representative images of Eph4 cells stained with **a,** phalloidin (yellow) and DAPI (cyan) **b,** anti-α-tubulin (yellow), Phalloidin (magenta) and DAPI (cyan) **c,** DAPI (cyan), Phalloidin (magenta), and expressing mAppple-Sec61β (yellow) at convex (upper panel) and concave curvature (lower panel), front of the cells at the edge is enlarged and zoomed, scale bar: 10µm. **d,** quantification of MDE for actin, microtubules, and ER at convex (green) and concave (pink) curvatures. n= 100, 90, 58, 61 from left to right. Data are mean ± s.e.m. ANOVA test.

**Supplementary figure 3. Curvature dependent response in mouse embryonic skin wounds. a,** mouse embryonic wounded skin tissue stained with phalloidin showing spontaneous concave (zoomed upper panel) convex (zoomed lower panel) curvature, the graph shows fluorescence intensity profiles of the solid lines, green dashed line shows convex curvature, Pink dashed lines show concave curvature **b,** quantification for the directionality of microtubules at convex and concave curvature in mouse embryonic skin wounds (n=26 at each curvature)

**Supplementary figure 4. Effect of blebbistatin and CK666. a,** Representative images of MDCK cells treated with DMSO (left) , CK666 (middle), and Blebbistatin (right) stained with phalloidin (grey), pink dashed lines mark concave curvatures and green dashed lines mark convex curvatures, scale bar: 10 µm. **b,** quantification for fraction of cells forming lamellipodia at convex or concave curvatures in cells treated with DMSO, CK666, or Blebbistatin (n=50). **c,** Representative images of MDCK cells expressing mApple-Sec61β (grey) treated with DMSO (left), CK666 (Middle), or Blebbistatin (right) in the middle of confluent monolayers, scale bar: 10 µm. **d,** Quantification of fraction of tubules in the cells in the middle of the monolayer treated with DMSO, CK666, or Blebbistatin; n= 59, 42, and 47 from left to right. Data are mean ± s.e.m, ANOVA test.

**Supplementary Figure 5. Computational models. a**, 3D visualization of the front portion of the cell indicating ER (green) embedded in the cytoplasm with actin cortex at the front edge (brown). **b,** Quantification of strain energy density during protrusion (left) and contraction (right) **c,** Schematic of cell with varying curvatures. **d,** Representative image showing the deformation of cell while experiencing protrusive force with ER sheets (left) and ER tubules (right). **e,** Representative image showing the deformation of cell while experiencing contractile force with ER sheets (left) and ER tubules (right).

**Supplementary figure 6. ER structure and microtubule alignment. a,** Representative images of MDCK cells expressing mApple-Sec61β (yellow) stained with anti-α-tubulin (magenta) and phalloidin (blue) migrating at convex (upper panel) and concave (lower panel) curvature.

**Supplementary figure 7. ER structural changes affect the mode of migration. a,** Representative super-resolution (SORA) images of cells expressing mApple-Sec61β (left), mCherry-Climp63 (middle) and Rtn4a-GFP (right); scale bar: 10μm. **b,** Representative images of MDCK WT (upper panel) and Climp63 (yellow, lower panel) migrating at convex curvature stained with phalloidin (magenta) and cortactin (green), scale bar: 5μm, fluorescence intensity line graphs for cortactin and phalloidin along the white lines marked, blue arrowhead shows cortactin peak at the edge of the cell **c,** Representative images of MDCK WT (upper panel) and Rtn4a (yellow, lower panel) migrating at concave curvature stained with phalloidin (magenta) and cortactin (green), scale bar: 5μm, fluorescence intensity line graphs for cortactin and phalloidin along the white lines marked, blue arrowhead shows cortactin peak at the edge of the cell

**G. Supplementary Video legends**

**Supplementary video 1.** LifeAct MDCK (Magenta) cell transfected with mApple-Sec61β (yellow) migrating at convex curvature. The movie is shown at 10 frames per second.

**Supplementary video 2.** LifeAct MDCK (Magenta) transfected cell with mApple-Sec61β (yellow) migrating at concave curvature. The movie is shown at 10 frames per second .

**Supplementary video 3.** LifeAct MDCK cell transfected with mApple-Sec61β (Black) migrating at convex curvature, zoomed at the migrating front show ER tubule dynamics. The movie is shown at 10 frames per second.

**Supplementary video 4.** LifeAct MDCK cell transfected with mApple-Sec61β (Black) migrating at concave curvature, zoomed at the migrating front show ER dynamics. The movie is shown at 10 frames per second.

**Supplementary video 5.** MDCK cells transfected with PA-Rac1 (Magenta) labeled with SiR-tubulin, first frame is before photoactivation. Yellow ROI depicts the region of photoactivation. The movie is shown at 10 frames per second.

**Supplementary video 6**. MDCK cells co-transfected with PA-Rac1 (Magenta) and GFP-Sec61β, first frame is before photoactivation. Yellow ROI depicts the region of photoactivation. The movie is shown at 10 frames per second.

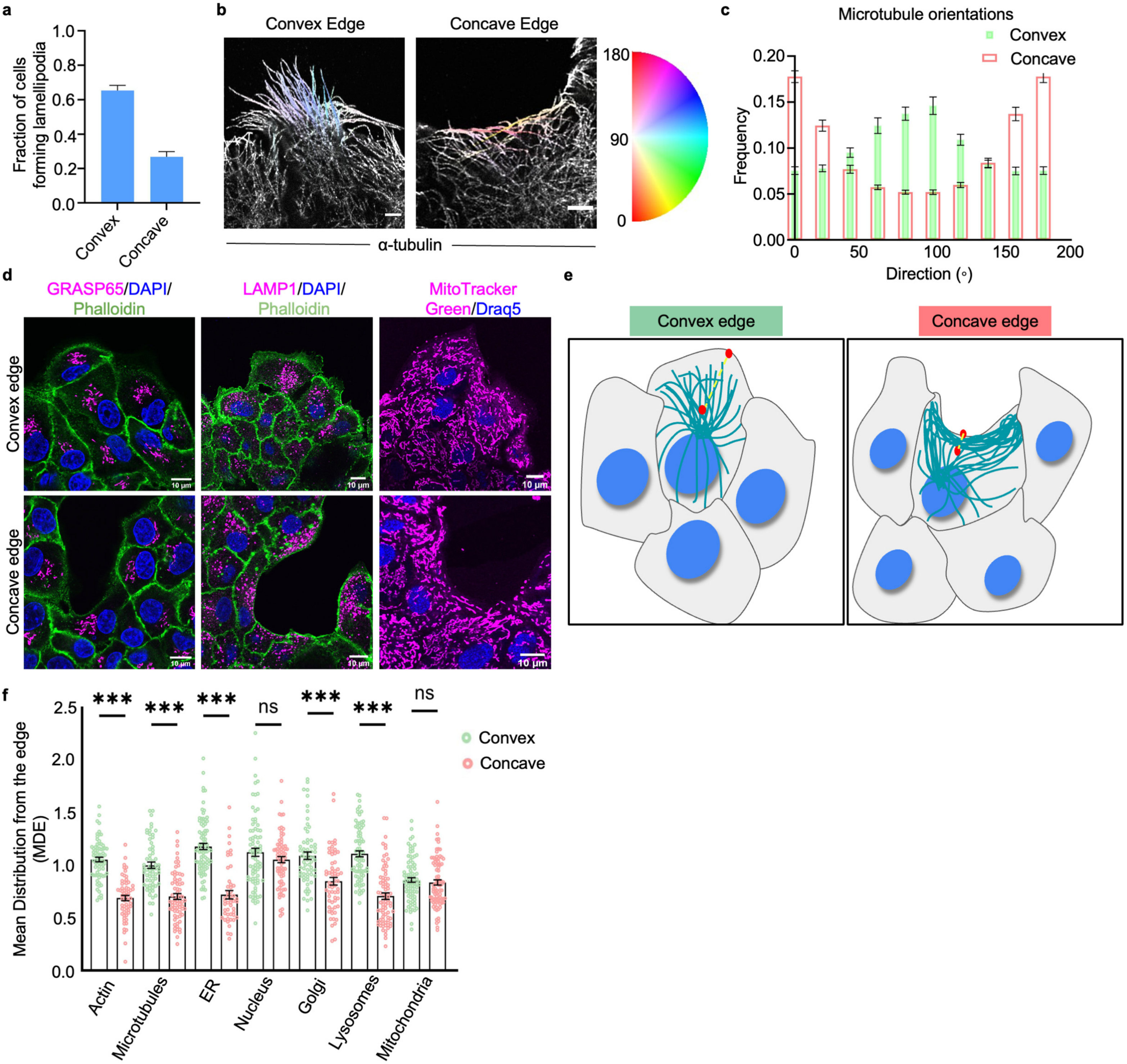


**Supplementary Figure 1**

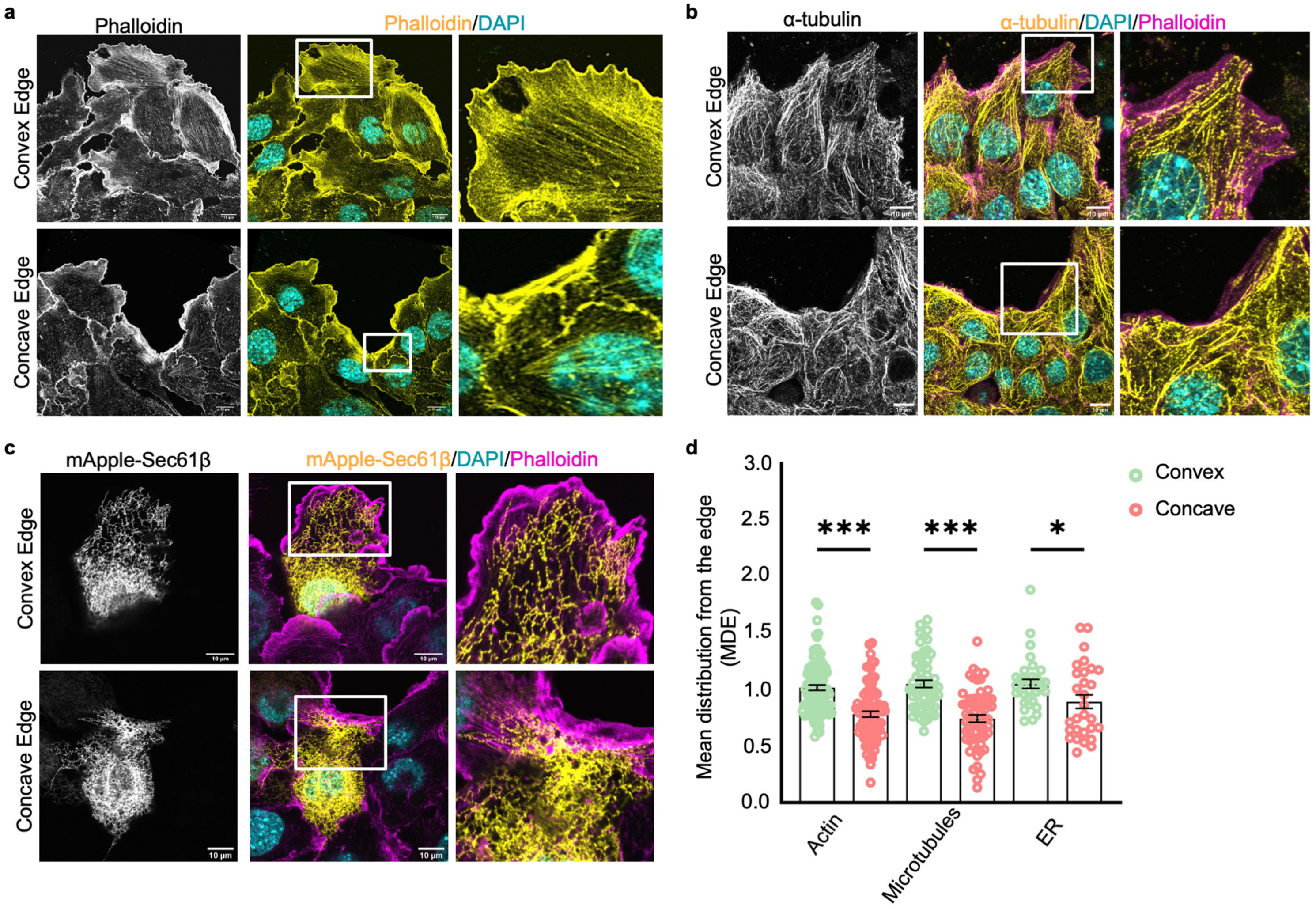


**Supplementary Figure 2**

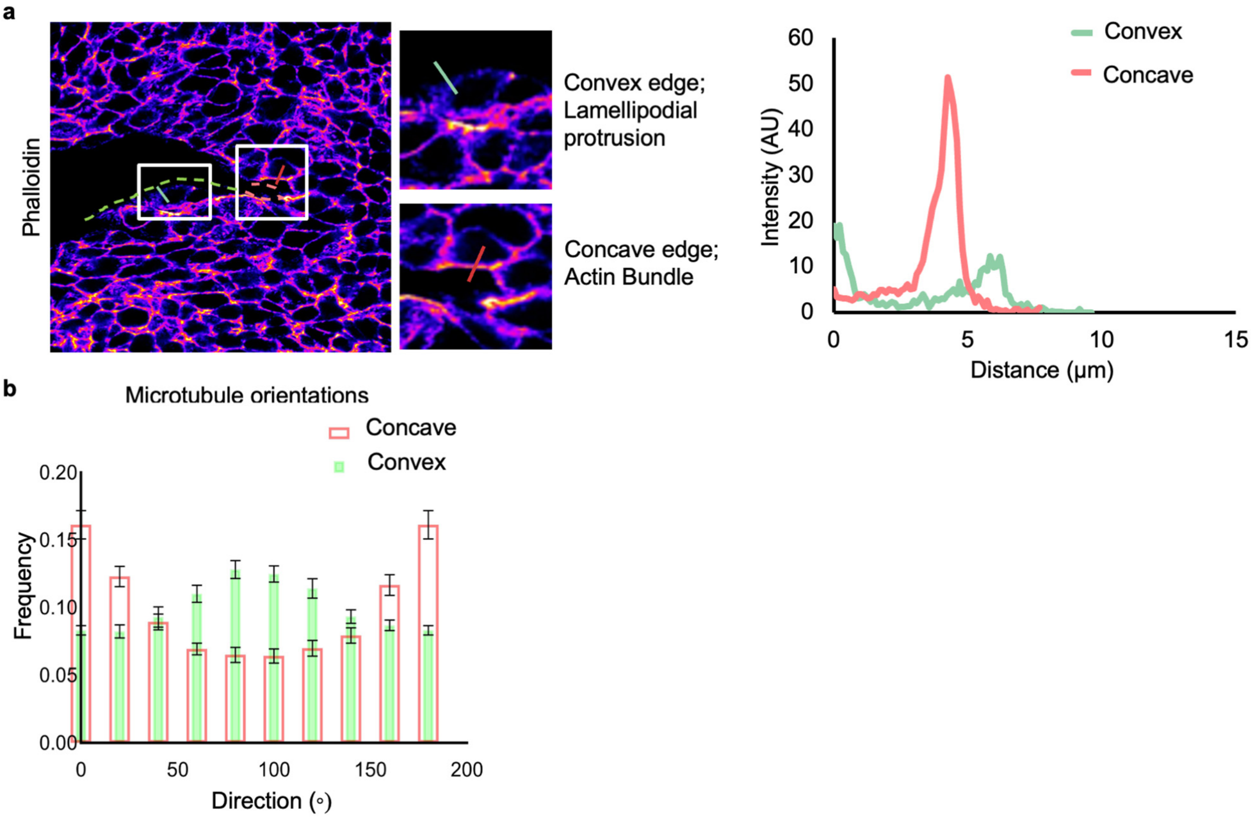


**Supplementary Figure 3**

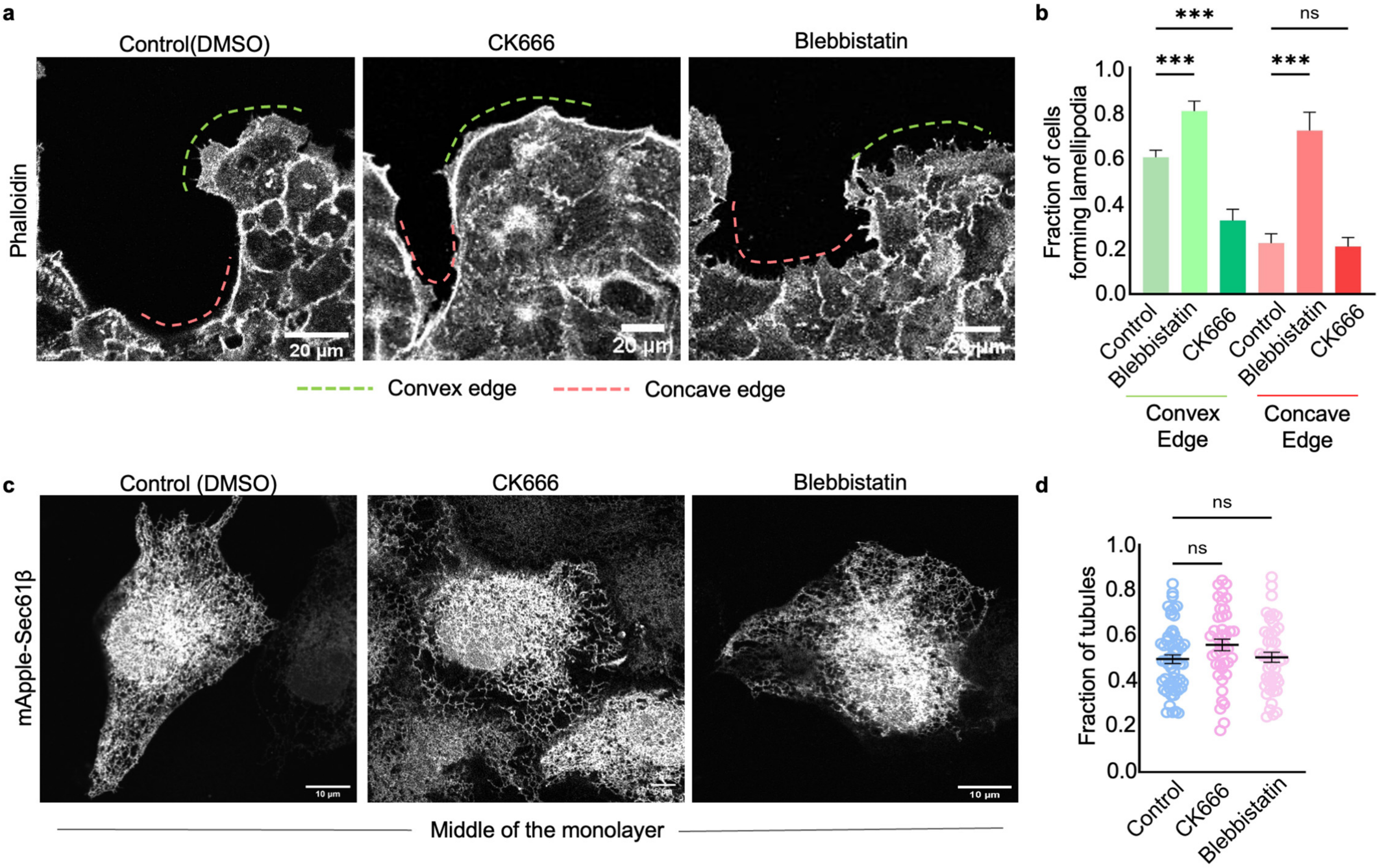


**Supplementary Figure 4**

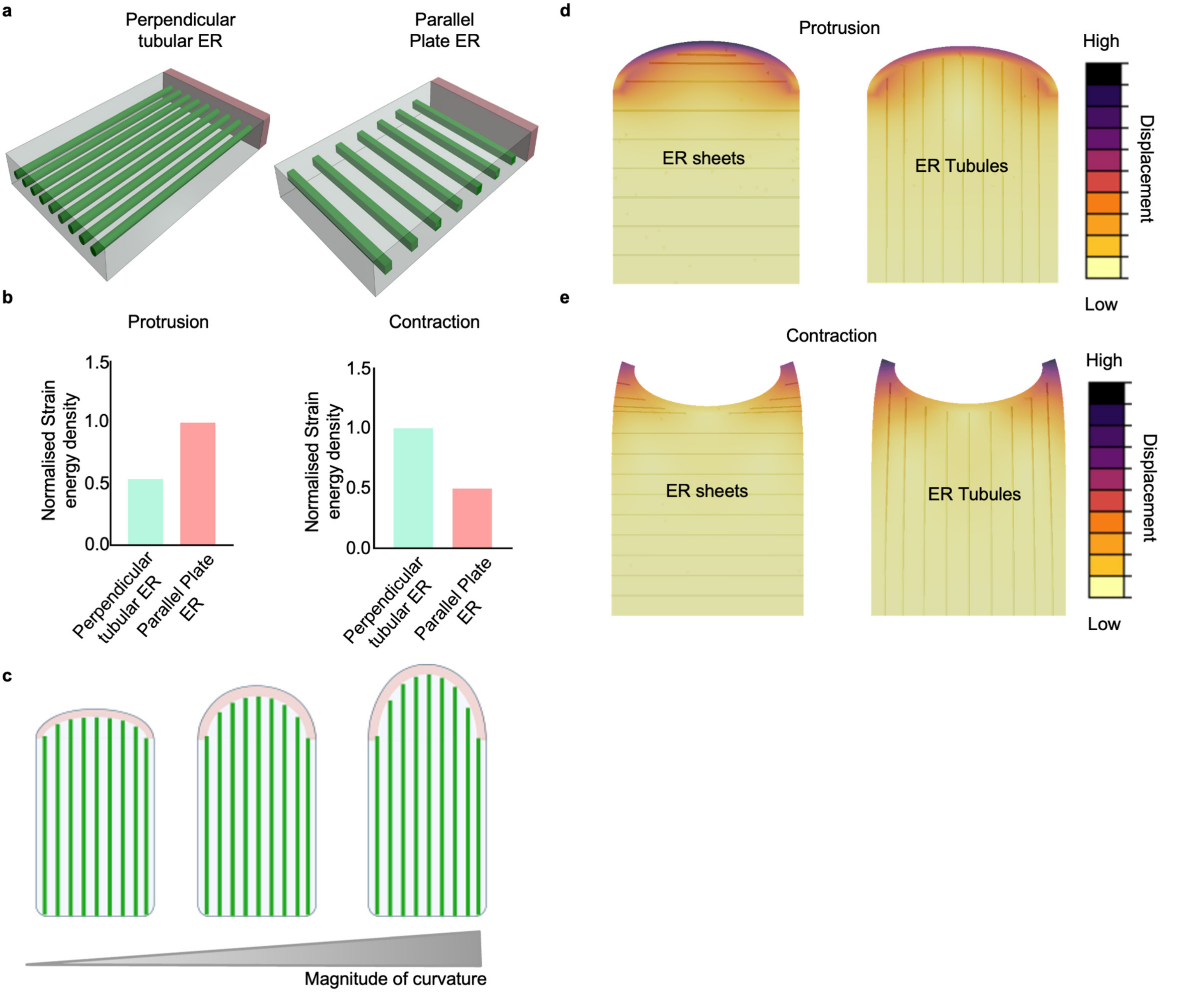


**Supplementary Figure 5**

a

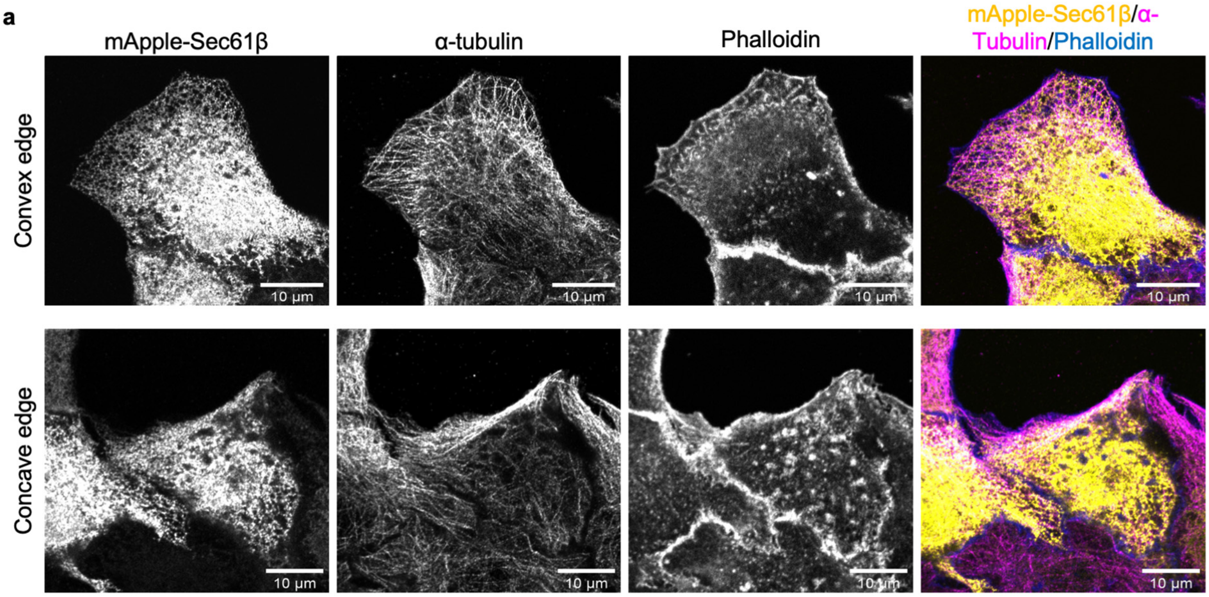


**Supplementary Figure 6**

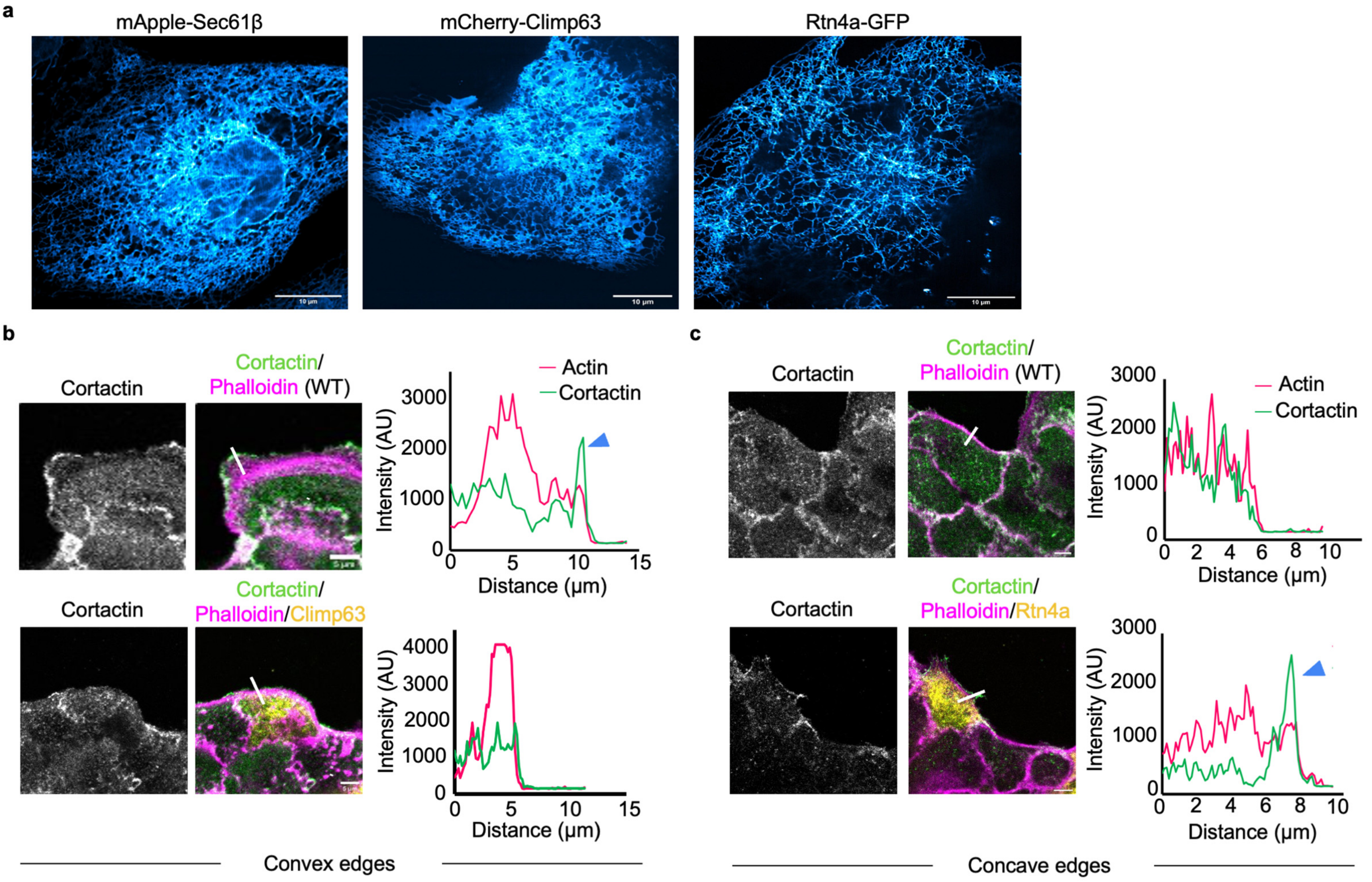


**Supplementary Figure 7**
